# Iron-mediated assembly of lactoferrin-alginate composites for iron encapsulation and structural stabilization

**DOI:** 10.64898/2026.04.21.719905

**Authors:** Yunan Huang, Tiantian Lin, Waritsara Khongkomolsakul, Jieying Li, Claire Elizabeth Noack, Younas Dadmohammadi, Alireza Abbaspourrad

## Abstract

Ternary composite systems formed by lactoferrin (LF), sodium alginate (Alg), and Fe(II) were designed to investigate their potential as an iron delivery platform with enhanced protein stability. The ternary LF-Alg-Fe (LAF) composites demonstrated distinct structures depending on the LF to Alg ratio and the Fe(II) concentrations. At an LF to Alg ratio of 8:2 and final Fe concentrations between 20-30 mM, the system formed complexes stabilized by electrostatic interactions. Whereas Alg-rich formulations formed hydrogels stabilized by Alg-Fe(II) egg-box cross-linking. Rheological analysis and swelling behavior indicated a higher mechanical strength in LF-rich complexes and stronger network integrity in Alg-rich hydrogels, while intermediate LF/Alg ratios showed weaker structures overall. Fourier-transform infrared spectroscopy (FTIR) spectra showed no changes in functional groups or polymer structures after composite formation, confirming composite formation via non-covalent interactions. Thermal studies indicated that these ternary systems improved LF stability, evidenced by preserved secondary structure after heating using circular dichroism (CD), and an increased denaturation temperature compared with free LF in differential scanning calorimetry (DSC). In addition, in LF-rich formulations the Fe(II) release in aqueous solution was ∼50% while in Alg-rich formulations it was much lower (< 10%). LF-Alg-Fe composites exhibit distinct structures governed by protein-polysaccharide interactions and iron-mediated cross-linking, providing a potential strategy for protein stabilization and iron fortification in food systems.

## 1 Introduction

Iron deficiency remains one of the most pervasive global health issues, affecting almost a quarter of the global population, especially among infants, young children, pregnant women, and individuals with chronic diseases (Breymann, 2015; Theurl et al., 2009). Despite the widespread implementation of iron supplementation and fortification programs, the incorporation of iron into delivery systems presents several technological challenges. Iron salts are highly reactive and can catalyze oxidative reactions, induce undesirable color changes, and generate off-flavors that negatively affect product quality and consumer acceptance (Kumari & Chauhan, 2022). In addition, interactions between iron and other food components may influence its stability and bioavailability during processing and storage (Hurrell, 2002). These challenges have motivated increasing research efforts toward developing delivery systems capable of stabilizing iron while maintaining its nutritional functionality.

To address the high reactivity of iron in food matrices, various delivery and encapsulation strategies have been explored to improve iron stability during processing and storage (Hurrell, 2022; Kumari & Chauhan, 2022). Biopolymer-based systems have proven to be promising because they can physically confine iron and modulate its local chemical environment. In particular, proteins and polysaccharides are widely used due to their biocompatibility and their ability to interact with metal ions (Gholam Jamshidi et al., 2024; Noack et al., 2025; Shen et al., 2017). Protein-polysaccharide complexes can form organized microstructures through electrostatic interactions, providing protective matrices that limit undesirable reactions between iron and other food components while maintaining its bioavailability.

Lactoferrin (LF) is an iron-binding glycoprotein present in milk and other secretory fluids, which can bind two Fe(III) ions per molecule (Rosa et al., 2017). Studies have shown that LF can improve iron bioavailability and enhance iron absorption, particularly in the distal gut where the pH is higher and iron is less soluble (Bokkhim et al., 2013; Mikulic et al., 2020). In addition to its nutritional relevance, LF is also known for its antimicrobial, antiviral, and anti-inflammatory properties, making it an attractive functional ingredient in food and nutraceutical applications (Hong et al., 2024; Jenssen & Hancock, 2009; Liu et al., 2018). However, LF is sensitive to thermal processing and can undergo structural changes at elevated temperatures (Bengoechea et al., 2011). The denaturation of LF not only reduces its functional properties but also diminishes its ability to bind and deliver iron effectively. Therefore, preserving the structural integrity of LF after thermal treatment is crucial to expand its application in iron fortification and other nutraceutical formulations.

Sodium alginate (Alg), a natural anionic polysaccharide derived from brown seaweed, is widely used in the food industry for its ability to form clean-label gels. The negatively charged carboxyl groups of alginate coordinate with divalent cations, forming ionically cross-linked egg-box structures (Cofelice et al., 2023; Dash et al., 2023; Hu et al., 2021). These egg-box units are further connected through hydrogen bonding, forming a three-dimensional hydrogel network.(Hu et al., 2021). Several studies have demonstrated that Alg enhances the thermal stability of LF through electrostatic interactions while retaining its iron-binding capacity, making LF-Alg systems a promising carrier for iron delivery (Bastos et al., 2018; Bokkhim et al., 2015). The combination of protein-polysaccharide interactions and ion-mediated crosslinking offers opportunities to design hybrid systems with tunable structures and improved stability for iron delivery. Iron may play a dual role in such systems, acting both as a micronutrient and as a coordination cross-linker that contributes to network formation. However, the structural organization and stability of ternary systems composed of LF, Alg, and iron remain poorly understood.

Based on the reported stability enhancement between LF and Alg, as well as the cross-linking property of iron, we hypothesized that the electrostatic interactions between LF and Alg would synergistically facilitate the formation of an LF-Alg-Fe (LAF) ternary composite system that would help protect LF from thermal denaturation and provide an Fe(II) delivery method. We expect such structures may provide a protective environment for LF while enabling iron encapsulation within the network.

In this study, we reported the formation of ternary LAF composites and their structure-dependent properties. The effects of LF/Alg ratio and Fe(II) concentration on structural outcome were examined to determine the conditions leading to either electrostatic complex formation or hydrogel network formation. Further, the rheological behavior, swelling properties, and microstructure of the composites were characterized to evaluate their structural stability. Finally, the thermal stability of LF within the composites and the retention of iron in aqueous environments were investigated to assess the potential of this system as a protein-stabilizing iron delivery platform.

## 2 Materials and methods

### 2.1 Materials

Bovine LF (Bioferrin 2000; Iron content >15 mg/100 g) was obtained from Glanbia Nationals, Inc. (Fitchburg, WI, USA). Food grade sodium alginate was purchased from Modernist Pantry, LLC (Eliot, ME, USA). Reagent grade ferrous sulfate heptahydrate, sodium chloride, sodium dodecyl sulfate, and L-ascorbic acid were purchased from Sigma-Aldrich (St. Louis, MO, USA). Reagent grade sodium hydroxide, hydrochloric acid, and urea were purchased from Fisher Scientific (Hampton, NH, USA). Ammonium acetate was purchased from Research Products International (Mt Prospect, IL, USA). Ferrozine (≥ 98%) was purchased from Cayman Chemical Company (Ann Arbor, MI, USA). All chemicals were of analytical or ACS reagent grade and used as received unless otherwise specified. All aqueous solutions were prepared using Milli-Q water (18.2 MΩ·cm).

### 2.2 Formation of LF-Alg-Fe (LAF) composite

The LAF composites were formed by an in-situ assembly method. LF and Alg stock solutions (20 mg mL^−1^) were prepared separately in Milli-Q water, with LF stirred magnetically at 25 °C for 30 minutes and Alg at 50 °C for 2 hours and then stored at 4 °C overnight to allow full hydration. An FeSO_4_ stock solution (1 M) was freshly prepared before use to avoid iron oxidation. All stock solutions were allowed to reach room temperature before use. For composite formation, the FeSO_4_ solution was added to the LF stock solution first, then the LF-Fe mixture and the Alg solutions were mixed at different volume ratios to achieve LF-Alg ratios 8:2 to 2:8 with final Fe concentrations between 0 and 30 mM. The mixtures were vortexed for 5-10 seconds, allowed to stand at room temperature for 30 min and stored at 4 °C for complete network development prior to analysis. After equilibration, one portion of the samples was used directly for characterization in hydrated form, while the remaining portion was freeze-dried for further characterization.

### 2.3 Characterization of LAF composites

#### 2.3.1 Zeta-potential

The zeta-potential was measured by the Zetasizer (Malvern Nano-ZS90, U.K) under the Smoluchowski mode at 150 V and 25 °C with disposable folded capillary cells used for measurement. All samples were diluted to 2 mg mL^−1^ with milli-Q water before measurement for better dispersion. All measurements were performed in triplicate, with a minimum of 11 runs per measurement.

#### 2.3.2 Rheological properties

The rheological properties of LAF composites were measured by the TA DHR3 rheometer (TA instrument, DE, USA) with the Peltier temperature control system. A 40 mm parallel plate geometry was used with the gap 1000 μm, and the temperature was fixed at 25 °C for all measurements. The flow behavior was measured under steady shear mode with the shear rate ranging from 0.01 to 100 s^-1^.

To determine the linear viscoelastic region (LVR), a strain sweep was first carried out for each sample at fixed frequency 1 Hz, within the strain range 0.01%-100%. Frequency sweeps were then performed under a constant shear strain within the LVR of all samples (0.1% for LAF composites), and the frequency range was set to 0.1-100 Hz. The Power Law model was used to fit the storage modulus (G’, Pa) and loss modulus (G’’, Pa) with frequency with the following equations:

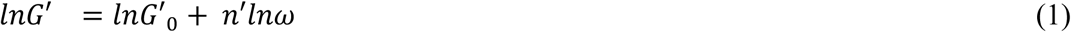

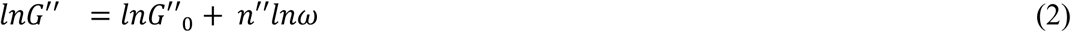

where G’_0_ and G’’_0_ are related to the strength of the network: a higher G’₀ value reflects increased material rigidity, whereas a higher G’’₀ suggests a greater viscous response, and n’, n” provide insight into the viscoelastic behavior.

#### 2.3.3 Swelling degree and Fe(II) diffusion

Equal masses of aqueous LAF composites were cast into silicone molds and freeze-dried to obtain uniformly shaped samples for standardized swelling and Fe diffusion measurements. The dried samples were then rehydrated at 25 °C, in milli-Q water or buffers at different pH levels for 30 min, after which the unabsorbed solution was removed by filtration through a Whatman qualitative filter paper (Grade 1, pore size 11 μm). The weight of LAF composites before and after swelling was measured and the swelling degree (SD) was calculated by the flowing equation:

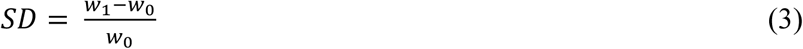

where w_0_ and w_1_ are the weight of the LAF composites before and after hydration, respectively.

The Fe(II) that diffused into the filtrate from the LAF composites was measured by ferrozine method (Huang et al., 2024) and standard curves for Fe(II) (0-0.5 mM) were established. The Fe(II) release was then calculated with the following equation:

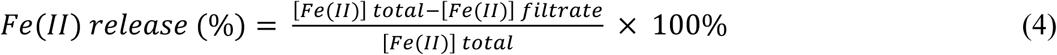

Where [Fe(II)] total is the concentration if all the Fe(II) was released from the LAF composite and [Fe(II)] filtrate was measured concentration in filtrate.

#### 2.3.4 Fourier transform infrared (FT-IR) spectroscopy

The FT-IR spectra of the starting polymers and powdered freeze-dried LAF composites were collected by IR spectrophotometer (Shimadzu IRAffinity-1S, Shimadzu Corp., Japan) between 400 to 4000 cm^−1^ with a resolution of 4 cm^−1^. A background correction was performed before each sample measurement, with 64 scans per measurement.

#### 2.3.5 Electrophoresis analysis

The presence of lactoferrin (LF) in redispersed LAF composites was analyzed by sodium dodecyl sulfate-polyacrylamide gel electrophoresis (SDS-PAGE) using a vertical mini gel electrophoresis system (Mini-PROTEAN Tetra Cell, Bio-Rad, USA). Polyacrylamide gels were prepared using a TGX FastCast Acrylamide Starter Kit (Bio-Rad, USA), and electrophoresis was performed using Tris-glycine-SDS running buffer.

Rehydrated LAF composites (2 mg mL^−1^ in phosphate buffer, 10 mM, pH 7.0) were mixed with a 2× Laemmli loading sample buffer at 1:1 volume ratio and heated in boiling water for 5 min. Subsequently, 20 μL of each sample was loaded onto the gel. Electrophoresis was carried out at 90 V for the stacking gel and 120 V for the separating gel.

After electrophoresis, the gels were stained with 0.15 w/v% Coomassie Brilliant Blue R-250 solution containing 50 v/v% methanol and 10 v/v% acetic acid for 30 min. The gels were then destained in a solution containing 10 v/v% methanol and 10 v/v% acetic acid. The destaining solution was replaced as needed until a transparent background was obtained.

#### 2.3.6 Scanning electron microscope (SEM)

The microstructures of LF and LAF composites were analyzed using a scanning electron microscope (SEM, Zeiss Gemini 500, Jena, Germany). A small amount of LF powder or freeze-dried LAF composites were applied to a pin stub with carbon tape and carbon coated before scanning, and a high efficiency secondary electron detector with a 20.0 μm aperture was used for scanning and photographed with the HE-SE2 signal (Lin et al., 2023). The accelerating voltage was 1 kV for imaging.

### 2.4 Thermal stability of LAF composites

#### 2.4.1 Differential scanning calorimetry (DSC) analysis

The thermal behavior of the LF and freeze-dried powdered LAF composites were analyzed using a modulated differential scanning calorimeter (MDSC 2910, TA instrument, USA).

Approximately 3 mg of each powdered sample was weighed into a standard aluminum pan, followed by the addition of 10 μL of Milli-Q water before sealing. The samples were allowed to sit overnight before measurement. Empty sealed aluminum pans were put in the heating chamber together with samples for reference. The heating program was set to scan from 30 to 160 °C at a rate of 10 °C/min. Thermal transition parameters, including the onset temperature (T_onset_), peak temperature (T_m_), and enthalpy change (ΔH) were determined with the Universal Analysis 2000 software (TA instruments).

#### 2.4.2 Thermal treatment of redispersed LAF composites

The stability of LAF composites during thermal treatment was evaluated by redispersing the freeze-dried powders in 2 mg mL^−1^ in phosphate buffer (10 mM, pH 7.0) and loaded into Pyrex-glass tubes. The suspensions (2 mL) were then heated at 95 °C in a water bath for 5 min and immediately placed in an ice bath to cool to room temperature.

The degree of Fe(II) oxidation, the presence of LF and its secondary structure were assessed for all samples before and after heating. The total Fe and Fe(II) concentration was evaluated with ferrozine method as described and the Fe(II) oxidation (%) was calculated by the following equation:

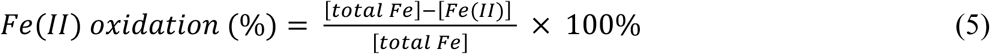

where the [total Fe] is the total measured iron concentration after reduction, and [Fe(II)] is the Fe(II) concentration measured directly in the sample.

#### 2.4.3 Circular dichroism (CD) spectra

The CD spectra of protein in LAF solutions were collected before and after thermal treatment to evaluate the protein secondary structure change during heating with JASCO-1500 CD Spectrometer (JASCO, Japan). To avoid signal gain and for better comparison, all samples were diluted to protein (LF) concentration 0.04 mg mL^−1^ before measurement. The scan range was set to 190-260 nm, and measurements were conducted using quartz cells with a 10 mm path length. The protein secondary structure was analyzed using the BeStSel web server (Miles et al., 2022) based on the spectra.

### 2.5 Statistical analysis

All measurements were carried out in duplicates or triplicates, and results were reported as mean ± standard deviations. Statistical analysis was conducted using JMP pro 19 software by one way ANOVA analysis and the difference between means values was evaluated by Tukey test (*p* < 0.05) .

## 3. Results and discussion

### 3.1 Formation mechanism of LAF

To determine the roles of both the Fe(II) concentration and LF/Alg ratio in determining the structure of the ternary system, samples were prepared across Fe(II) concentrations from 0 to 30 mM and LF/Alg ratios from 8:2 to 2:8. The resulting materials were evaluated first by visual observation (tube inversion) and then by their zeta potential (**Fig. 1**).

**Fig. 1.**
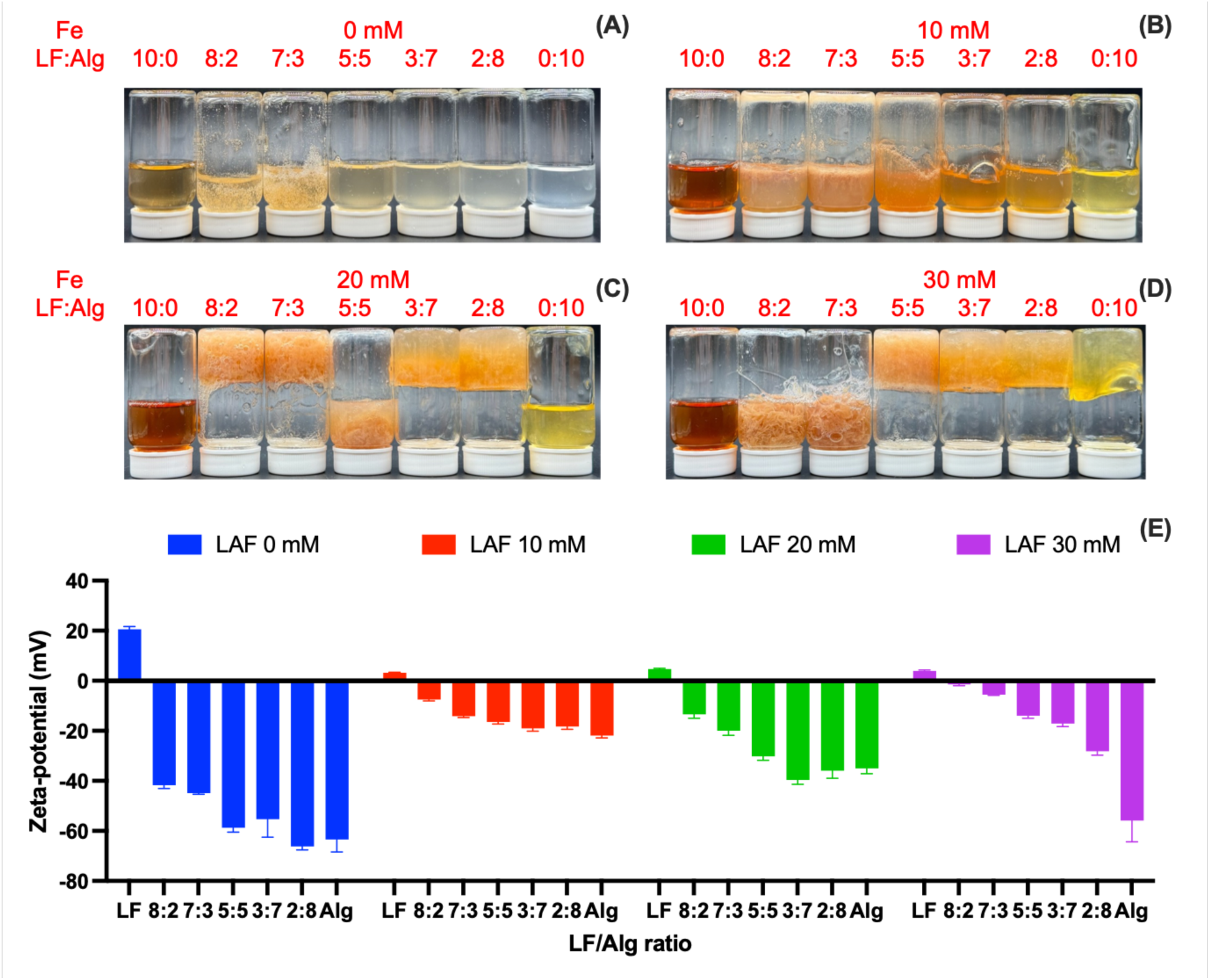
LAF formation condition with different LF-Alg ratios and Fe concentrations (A∼D: 0∼30 mM Fe concentration in final hydrogel) and the corresponding zeta potential (E).

In the absence of Fe(II) and at low Fe(II) concentrations (≤10 mM), no stable hydrogels were observed (**Fig. 1A-B**). Mixtures with LF-Alg ratios of 8:2 and 7:3 become viscous (0 mM Fe(II)) or turbid (10 mM Fe(II)), suggesting partial electrostatic complexation between LF and Alg. The zeta potential of LF-Alg mixtures without Fe(II) remained highly negative (**Fig. 1E**), indicating that the weak electrostatic interactions alone were insufficient to support the formation of a gel network.

As the Fe(II) concentration increased to 20 mM, stable hydrogels were observed at high LF ratios (LAF 8:2 and 7:3, **Fig. 1C**), accompanied by less negative zeta potential values indicating partial charge neutralization between LF and Alg (**Fig. 1E**), complemented by Fe(II)-mediated cross-linking. At higher Alg ratios, the zeta-potential remained strongly negative, indicating weaker electrostatic interaction (**Fig. 1E**). Nevertheless, gel formation was still observed, which suggested the Alg-Fe cross-linking allowed the network formation. However, for the sample with LF to Alg ratio 5:5 and the Fe(II) concentration 20 mM, neither electrostatic complexation nor Fe-mediated cross-linking was sufficiently strong enough to support a stable hydrogel structure.

Increasing the Fe(II) concentration to 30 mM, the formation of LAF hydrogels was enhanced at high Alg to LF ratios (LAF 5:5 to 0:10) due to higher degree of Fe cross-linking. However, at LF-rich ratios (8:2 and 7:3), the systems no longer formed stable hydrogels but instead produced large flocculates and complexes (**Fig. 1D**). This transition, caused by increasing the Fe(II) concentration from 20 to 30 mM, is consistent with the much lower zeta potential values measured at LF to Alg ratios of 8:2 and 7:3 **(Fig. 1E)**, indicating near charge neutralization and stronger electrostatic complexation between LF and Alg. The increased positive charge contributed by Fe(II) likely promoted the formation of ternary complexes, resulting in a structural transition from hydrogels to complex aggregates.

Under very low concentration conditions (0.2% total solids, LF/Alg 8:2-5:5, Fe(II) 3 mM, **Fig S2**), LF/Alg mixtures were not self-supporting upon inversion, but did show measurable turbidity, indicating the formation of electrostatic complexes. Upon the addition of NaCl, the turbidity of these complexes decreased progressively as the sodium and chloride ions surrounded the LF and Alg and neutralized their opposing charges, thus supporting that these complexes are primarily stabilized by electrostatic interactions.

To further verify the interaction mechanisms, two representative systems (LAF 8:2 30 mM and LAF 2:8 30 mM) were selected for blocking experiments using NaCl and urea to disrupt electrostatic and hydrogen-bonding interactions. For the LAF complex with an LF to Alg ratio of 8:2 and an Fe(II) concentration of 30 mM, where electrostatic complexation dominated, the addition of NaCl eliminated flocculation and produced a clear solution and urea had little effect (**Fig. S1A-B)**. In contrast, the LAF hydrogel with a LF to Alg ratio of 2:8 and an Fe(II) concentration of 30 mM, which formed a robust three-dimensional hydrogel network due to Alg-Fe egg-box cross-linking, when urea was added the hydrogel structure was disrupted, and NaCl had only limited weakening effect at high concentration (1 M) (**Fig. S1D**), suggesting that although Na^+^ can compete with Fe(II) for alginate binding sites, it is not sufficient to fully disrupt the network.

This behavior is likely related to the relatively strong Fe-Alg coordination as well as the additional stabilization provided by LF within the ternary system. In contrast, Fe(II) alone does not form robust alginate hydrogels. Together, these results suggest that the stability of the LAF hydrogels arises from the combined effects of Fe-mediated cross-linking and LF-Alg interactions, rather than Fe(II) cross-linking alone

Because stable hydrogel structures were only obtained at Fe(II) concentrations of 20 and 30 mM, the ternary systems formed at these two Fe(II) concentrations were selected for subsequent characterization and structural analysis. The effect of pH on LAF hydrogel formation demonstrated the role of the electrostatic interaction between Alg-LF. The initial pH of the mixtures was approximately 5.8-5.9, and additional experiments were done at pH 5.0 and 6.0 (**Fig. S3**). Alg has a relatively low pKa (∼3.5) and is therefore largely deprotonated under the experimental pH conditions investigated here, resulting in consistently negative zeta potential values. LF has a lower positive charge at lower pH conditions (pH 5.0), specifically, the LF to Alg ratio 8:2 30 mM sample had a near-neutral zeta-potential compared to the system at pH 6. This change in charge with pH, reduced electrostatic repulsion and promoted the formation of dense complexes. This resulted in more pronounced flocculation and precipitation-like behavior rather than continuous gel networks (**Fig. S3E-F**).

As the pH increased to 6.0, the overall zeta potential shifted to more negative values, indicating reduced net positive charge on LF and weaker LF-Alg electrostatic interactions. Under these conditions, hydrogel formation was weakened at high Alg ratios, particularly at 20 mM Fe(II). In LF-rich systems, the increased negative charge suppressed excessive aggregation; for example, at LF/Alg 8:2 and 30 mM Fe(II), although some flocculated structures were still observed, the system predominantly exhibited a gel-like morphology rather than forming fully precipitated complexes (**Fig. S3C-D**).

Overall, these results suggest that pH modulates the balance between electrostatic complexation and network formation through its influence on zeta potential. At lower pH, near-neutral charge conditions favor complex formation in LF-rich systems, whereas at slightly higher pH, increased electrostatic repulsion supports the formation of more continuous hydrogel structures. However, the overall differences observed across the tested pH range were not sufficiently pronounced to alter the main structural trends. Therefore, subsequent experiments were conducted at the unadjusted pH (∼5.8–5.9).

### 3.2 Characterization

#### 3.2.1 Rheology Steady flow behavior

The flow behavior of LAF composites evaluated the effect of shear rate on viscosity (**Fig. 2**). The apparent viscosity (*η_a_*) of all samples decreased with increasing shear rate (*γ*), indicating a shear-thinning behavior. This was a characteristic of most non-Newtonian fluids formed by biopolymers and attributed to the orientation and disentanglement of polymer chains under shear (Lin & Fernández-Fraguas, 2020).

**Fig. 2.**
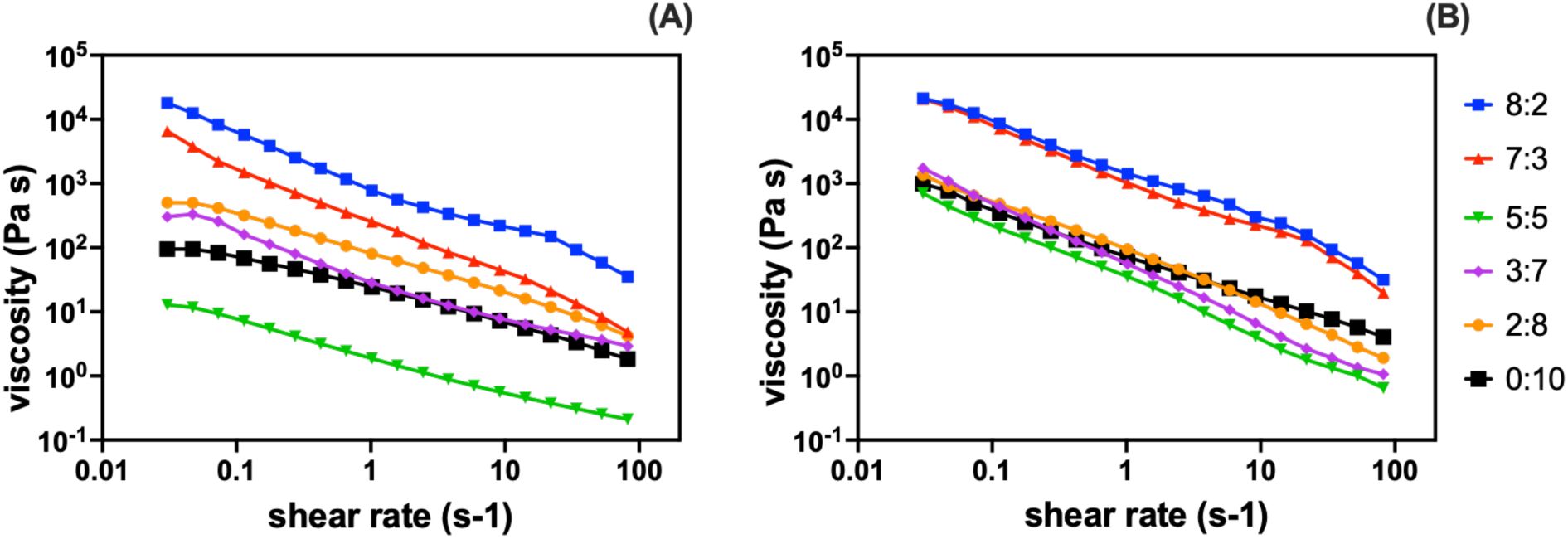
Flow behavior of LAF with different LF-Alg ratio and Fe concentrations at 25° C in the steady flow test. (A: 20 mM Fe; B: 30 mM Fe)

LAF complex systems with LF-rich compositions (8:2 and 7:3) exhibited higher viscosities than other ratios at both Fe(II) concentrations (20 and 30 mM), indicating stronger interactions within the ternary system (**Fig. 2**). In particular, LAF complexes at LF to Alg ratios of 8:2 showed slightly higher viscosity than that of 7:3, especially at 20 mM Fe(II), which is consistent with their lower zeta potential (**Fig. 1E)** and suggests stronger LF-Alg electrostatic interactions.

As the Alg ratio increased, electrostatic interactions between LF and Alg gradually weakened, while Fe-Alg cross-linking became more dominant. This compositional shift resulted in a decrease in viscosity at intermediate ratios, followed by a recovery at higher Alg ratios. At 20 mM Fe(II) (**Fig. 2A**), LAF 5:5 had the lowest viscosity, indicating a weak network structure where neither electrostatic interactions nor Fe-mediated cross-linking were dominant. As the Alg ratio increased further, viscosity increased again, with LAF 2:8 20 mM Fe(II) hydrogel showing higher viscosity than LAF 3:7 20 mM Fe(II), suggesting increased Fe-Alg cross-linking. The binary Alg-Fe system (0:10) showed lower viscosity than the ternary systems, highlighting the synergistic contribution of LF-Alg electrostatic interactions to network stabilization.

Increasing Fe(II) concentration to 30 mM further strengthened the network structure (**Fig. 2B**). The LF-rich complexes (LF/Alg ratio 8:2 and 7:3) remained the most viscous, and their viscosity only increased slightly with higher Fe(II) concentration. This is because at LF-rich ratios, the system was already in a complex-dominated state at 20 mM Fe(II) (**Fig. 1C**). Rather than strengthening a continuous network, increasing the Fe(II) concentration further led to more dense, flocculated aggregates (with zeta potential approaching neutral, **Fig. 1D &E**), and therefore only resulted in a small increase in viscosity. In contrast, Alg-rich hydrogel samples showed a more pronounced increase in viscosity when the Fe(II) concentration increased, particularly for LF to Alg ratios of 5:5 and Fe(II) 30 mM, indicating that Fe-Alg cross-linking becomes increasingly important in stabilizing the network at higher iron concentrations. This contrast reflects the different structures of the two systems, with LF-rich formulations dominated by complexes and Alg-rich formulations forming more continuous network structures, resulting in their distinct viscosity responses.

### Viscoelastic behavior

The viscoelastic behavior of LAF hydrogels or complexes evaluated their network stability and structural strength. A strain sweep was done to determine the linear viscoelastic region (LVR) for all samples (**Fig 3A-B**). In strain sweep measurement, the storage modulus (G’) presents the elastic or solid-like behavior of the material, while the loss modulus (G’’) reflects the viscous or liquid-like component. At both Fe(II) concentrations (20 and 30 mM), G’ was higher than G’’ for all samples except for the mixture of Alg and Fe without LF, indicating a predominantly solid-like structure of both LAF complexes and hydrogels.

**Fig. 3.**
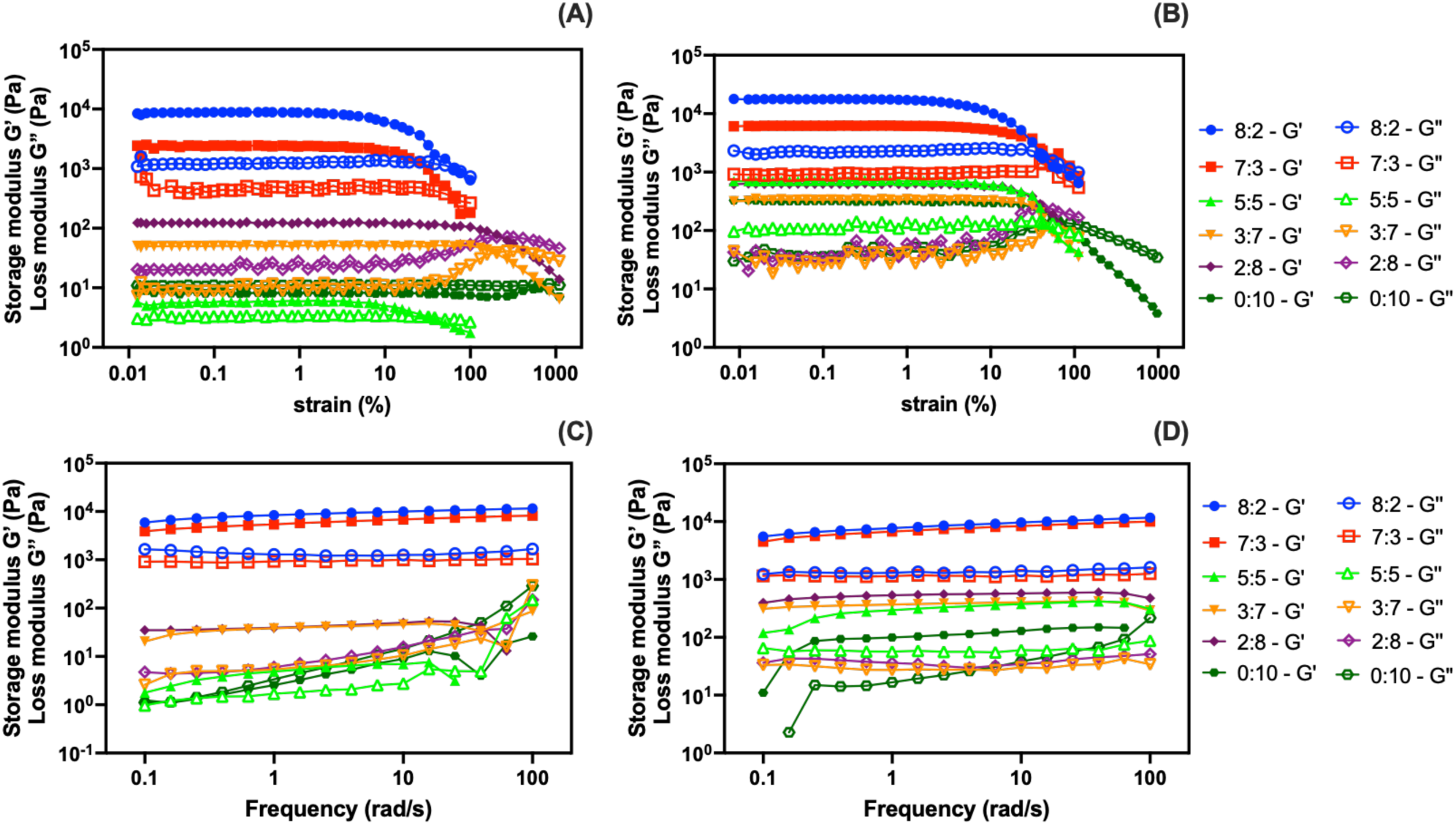
Strain sweep curves (A-B) and frequency sweep curves (C-D) of LAF with different LF-Alg ratio and Fe concentrations at 25° C. The strain sweeps were conducted at frequency 1 Hz, and the frequency was conducted at strain 0.1%. (A, C: 20 mM Fe; B, D: 30 mM Fe)

At Fe(II) 20 mM concentrations, the complexes formed at LF to Alg ratio of 8:2 showed the highest G’ and G’’, followed by the LF to Alg ratio of 7:3, reflecting rigid structures formed through strong LF-Alg electrostatic interactions. In contrast, LF to Alg ratio of 5:5 had the lowest G’ and the G’’ value as well as a smaller difference between G’_0_ and G’’_0_ indicating a weak network structure where neither electrostatic interactions nor Fe-mediated cross-linking were dominant. As the Alg concentration increased (LF to Alg ratios of 3:7 and 2:8), G’ and G’’ increased again due to stronger Alg-Fe interactions. The Alg-Fe binary mixture (no LF) showed a lower G’ and G’’ than most ternary systems due to the absence of electrostatic interactions, although it was stronger than the LF to Alg ratio of 5:5 at 20 mM of Fe(II) due to higher Alg ratio and stronger hydrogen bonding.

At 30 mM Fe(II) concentrations, a similar trend was observed, but with overall higher moduli. In particular, the G’ and G’’ values of the hydrogel with an LF to Alg ratio of 5:5 increased markedly and exceeded those of hydrogels with LF to Alg ratios of 3:7 and 2:8. We attribute this observation to a higher degree of Alg-Fe cross-linking which reinforced the network structure.

Consistent with this trend, the flow point, which is defined as the strain where G’ and G’’ intersect, was lowest for the LF to Alg ratio of 5:5, indicating the lowest resistance to deformation (**Fig. 3A-B**). LF-rich systems showed intermediate stability, whereas Alg-rich systems exhibited higher resistance to deformation due to increased cross-linking. The Alg-Fe binary system (no LF) showed G’’ > G’ across the entire strain range and therefore no flow point.

Frequency sweep measurements were then performed within the LVR at strain 0.1% (**Fig. 3C-D)**. For all samples, G’ was greater than G’’ (tan δ < 1) at low frequencies, indicating predominantly elastic behavior. With frequency increasing, at LF-rich ratios (8:2 and 7:3), the LAF complexes and hydrogels with 20 mM of Fe(II), G’’ increased gradually at >1 rad/s, showing a viscoelastic solid behavior where the network is stable at rest but easily deforms under increasing stress. While for both complexes and hydrogels with 30 mM Fe(II), the G’ and G’’ were almost parallel within the experimental range, which indicates weak gels as the samples maintained a stable network structure under sweep (Rao, 2010).

Quantitative analysis using the power law model (**Table S1**) further confirmed these trends. LF-rich systems (LF to Alg ratios of 8:2 and 7:3) showed the largest lnG’_0_-lnG’’_0_ values together with low n’ and n’’ (∼0.1), indicating rigid networks with minimal frequency dependence. As the Alg ratio increased, electrostatic interactions weakened while Alg-Fe cross-linking became more dominant. Increasing Fe(II) concentration to 30 mM further strengthened the networks, with higher lnG’_0_-lnG’’_0_ values observed across all compositions.

Overall, the rheological results indicate that the structure of LAF complexes and hydrogels is governed by two primary interactions: LF-Alg electrostatic complexation and Alg-Fe cross-linking. LF-rich systems (complexes dominated) exhibited higher rigidity due to stronger electrostatic interactions, whereas Alg-rich systems (hydrogel dominated) were strengthened by increased Alg-Fe cross-linking. At the intermediate ratio (LF to Alg ratio of 5:5), both interactions were relatively weak, resulting in lower overall gel strength. Nevertheless, this system remained stronger than the binary Alg-Fe mixture (0:10), highlighting the role of LF-Alg electrostatic interaction in stabilizing the ternary network.

#### 3.2.2 Swelling degree and Fe(II) release

Swelling behavior reflects hydrogel network strength and directly influences the Fe(II) release within the system in different environments (Li & Mooney, 2016). Understanding the swelling behavior will allow targeted Fe(II) release for delivery and sustained bioavailability in diverse food processing or physiological conditions.

The swelling degree (SD) of freeze-dried LAF composite samples was evaluated after immersion in Milli-Q water, pH 3.0 buffer, and pH 7.0 buffer for 30 min, and the photos of the LAF samples before and after swelling were captured (**Fig. 4A-B and S4-5**). Overall, the LAF complexes and hydrogels with 20 mM Fe(II) exhibited higher SD than those at30 mM Fe(II) (**Fig. 4A-B**). The increased Fe(II) concentration strengthened Alg-Fe cross-linking, resulting in denser networks with reduced water uptake capacity, and therefore the SD was lower.

**Fig. 4.**
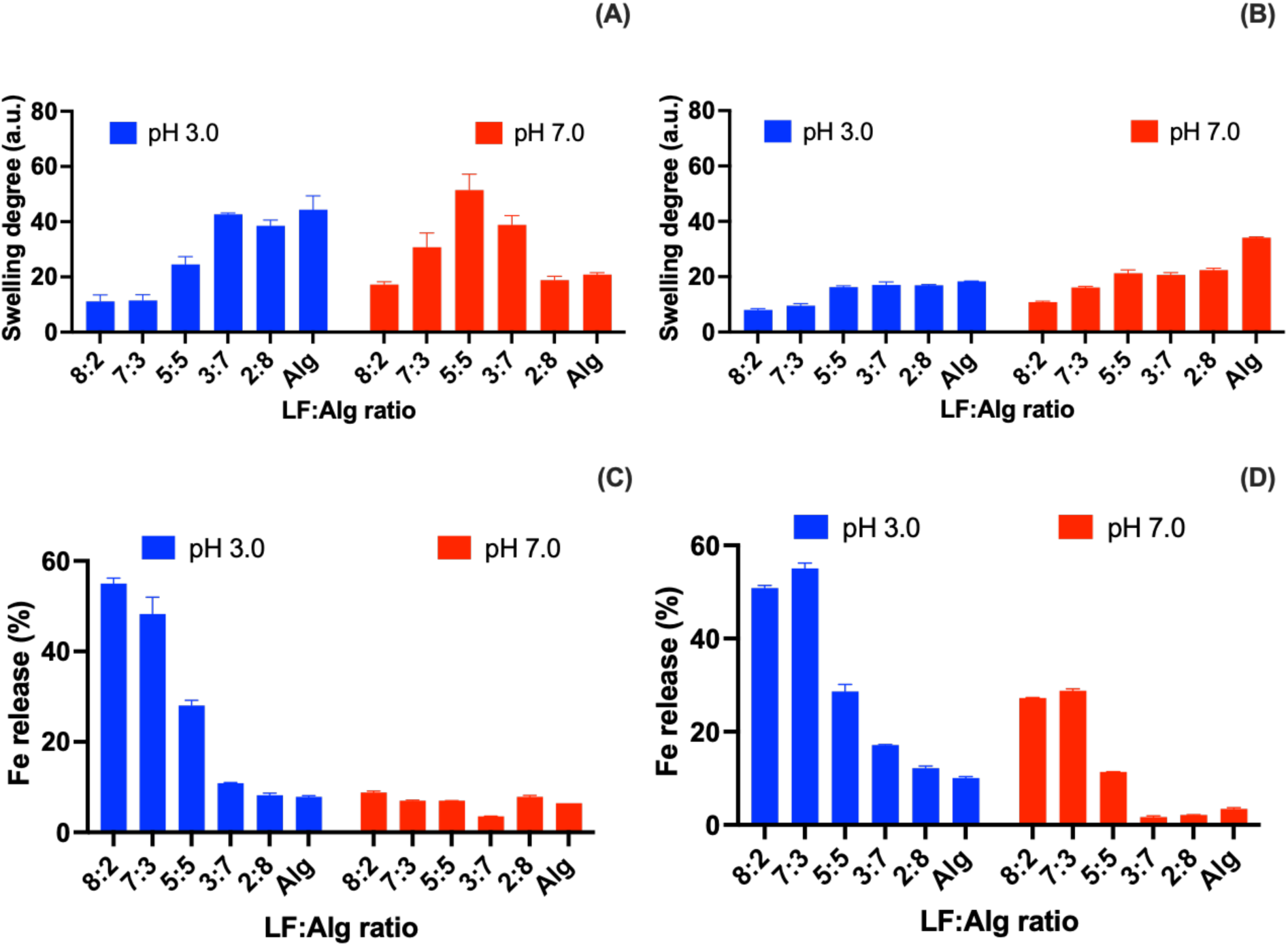
The swelling degree (A-B) and Fe release (C-D) of LAF with different LF-Alg ratios and Fe concentrations (A,C: 20 mM; B, D:30 mM).

The swelling behavior was strongly dependent on the LF to Alg ratio. LF-rich systems (LF:Alg 8:2 and 7:3) showed consistently lower SD across all conditions, and samples at these ratios with 30 mM Fe(II) did not maintain an intact hydrogel structure, but instead they appeared as aggregated particles (**Fig. S5B**), indicating the lack of a continuous 3D network. In contrast, Alg-rich systems (LF:Alg 5:5 to 0:10) remained stable hydrogel characteristics after swelling, with SD increasing as the Alg ratio increased, reflecting the formation of a cross-linked Alg-Fe network.

The effect of pH was more obvious for LAF composites with 20 mM Fe(II). Under acidic conditions, higher SD was observed compared to neutral conditions, particularly for Alg-rich samples (**Fig. 4A and S5A**). This can be attributed to protonation of Alg carboxyl groups, which weakens Fe-Alg interactions and reduces cross-linking density, allowing the network to swell more extensively(Li & Mooney, 2016). At neutral pH, stronger Alg-Fe cross-linking resulted in more compact structures and lower SD. Samples with no LF (LF:Alg 0:10) with 20 mM Fe(II) even exhibited reduced size and partial dissolution, likely due to increased Alg solubility upon carboxyl deprotonation at pH 7.0. In comparison, LAF complexes and hydrogels with 30 mM Fe(II) showed similar SD across all pH conditions, indicating that stronger cross-linking imparted by higher Fe(II) concentrations reduced pH sensitivity.

The Fe(II) release was measured by the concentration of Fe(II) that leached from the supernatant after swelling (**Fig. 4C-D**). Overall, Fe(II) release was governed by pH and LF to Alg ratio. Higher iron release was observed under acidic conditions, where protonation of Alg reduced available binding sites for Fe, weakening cross-linking and promoting iron release. In contrast, lower apparent Fe(II) release at neutral pH may be associated with reduced Fe(II) solubility under neutral conditions (Morgan & Lahav, 2007).

The LF to Alg ratio also influenced Fe(II) retention. LF-rich systems exhibited higher Fe(II) release due to weaker Fe(II) binding, as electrostatic complexes provide limited immobilization compared to cross-linked networks. In contrast, Alg-rich hydrogels retained Fe(II) more effectively, as Fe(II) was incorporated into the egg-box structure and physically confined within the network.

In Alg-rich hydrogels, increasing Fe(II) from 20 to 30 mM had limited impact on Fe(II) release, as Fe(II) was already strongly immobilized within the network. However, in LF-rich systems, higher Fe(II) concentration led to increased release, likely due to excess unbound or weakly associated Fe(II).

Overall, swelling and iron release depend on the balance between LF-Alg electrostatic interactions and Alg-Fe cross-linking, with cross-linked hydrogel structures providing enhanced iron retention and reduced sensitivity to pH levels.

#### 3.2.3 Fe oxidation of freeze-dried LAF

The oxidation state of the iron ions determines their bioavailability, with Fe(II) having higher absorption efficiency with fewer side effects compared with Fe(III) (Santiago, 2012). Therefore, the oxidation behavior of Fe ions within the ternary systems was evaluated (**Fig. 5)**.

**Fig. 5.**
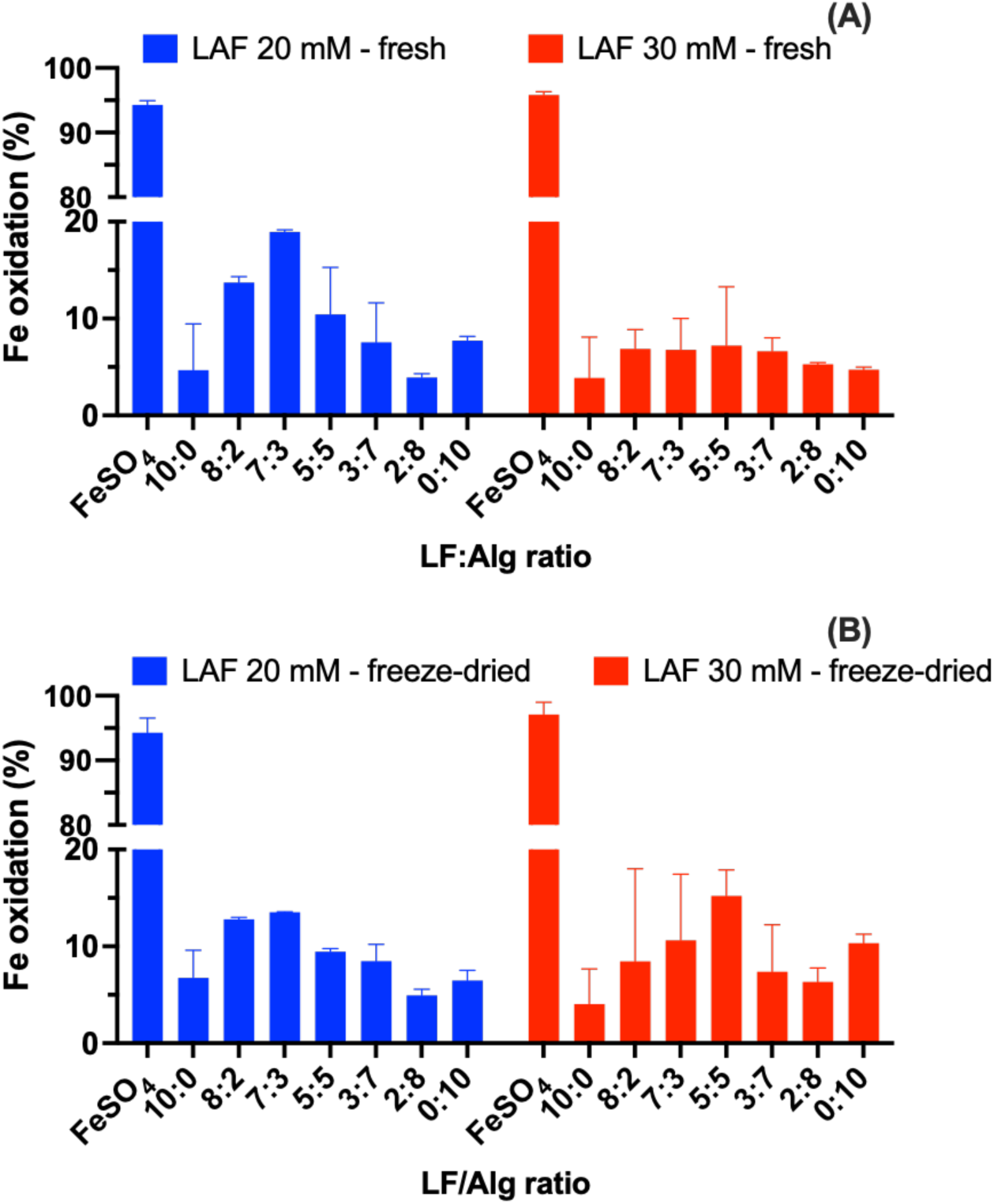
Fe oxidation of fresh (A) and freeze-dried (B) LAF at different LF-Alg ratios and Fe concentrations. The oxidation rates for all LAF 10:0-0:10 groups are not significantly different (ns, p>0.05).

Free FeSO_4_ at pH 5.8-5.9 showed pronounced instability in aqueous solution, with obvious precipitation (**Fig. S6C**) and rapid oxidation (> 90%, **Fig. 5A and S6D**, (Morgan & Lahav, 2007). In contrast, LF significantly improved Fe(II) stability, with LF only mixtures (LAF 10:0) maintaining low oxidation levels (< 10%) at both Fe(II) concentrations. This is attributed to the intrinsic antioxidant properties of LF (Dyrda-Terniuk & Pomastowski, 2023).

Incorporation of alginate enhanced Fe(II) stability. The formation of Alg-Fe cross-linked networks immobilized Fe(II), thereby limiting its accessibility to oxidizing species such as dissolved oxygen. As a result, freshly prepared LAF hydrogels followed by complexes showed substantially lower oxidation compared to free Fe(II), similar to the LF-only systems (**Fig. 5A**).

The extent of Fe(II) oxidation in LAF complexes and hydrogels was strongly dependent on network structure. LF to Alg ratios controlled the balance between electrostatic interactions and Alg-Fe cross-linking, leading to a non-monotonic trend in oxidation levels. Specifically, intermediate compositions such as LF to Alg ratios of 7:3 with 20 mM Fe(II) and LF to Alg ratios of 5:5 with 30 mM Fe(II), which formed relatively weaker networks, showed reduced protection against oxidation. While the higher Fe(II) concentration (30 mM) generally resulted in lower oxidation, likely due to enhanced Alg-Fe coordination and a more densely cross-linked network.

After freeze-drying, the LAF composite powders maintained oxidation levels comparable to freshly prepared samples (**Fig. 5B**), indicating that the encapsulation structure remained intact during dehydration. During storage at 4 °C for 120 days, oxidation in the LF only systems increased markedly (from ∼5% to >20%), whereas LF to Alg ratios of 8:2 to 0:10 exhibited less than a onefold increase (**Fig. S7**). This further confirms that Alg-Fe cross-linking provides sustained protection of Fe(II) against oxidation over prolonged storage.

#### 3.2.4 FT-IR

FTIR spectra of native biopolymers and freeze-dried LAF composites powders showed that the hydrogel and complexes formation did not change the functional groups present but did provide evidence of changes in bonding interactions for LF (**Fig. S8**). For LF, the amide I and amide II bands, which are characteristic of protein secondary structure, were observed at 1643, and 1511 cm^−1^, respectively. The free amino acid O-H stretching vibrations appeared at 3200-3300 cm^−1^ (Barth & Zscherp, 2002), while the band at 1039 cm^−1^ was associated with the C-N or C-C bond stretching vibration of sugar moieties (McAvan et al., 2020).

In the LAF complexes and hydrogel samples, the amide I vibration slightly right shifted, while the amide II intensity gradually decreased, suggesting that protein-Fe complexation altered the H-bond network. The band at 1039 cm^−1^ was also red shifted when Alg was added, which is consistent with reported sugar-related vibrations in the range of 1035-1055 cm^−1^ (Lin et al., 2022a). The intensity of the band at 1040 cm^−1^ increased after adding Fe(II) is attributed to enhanced C-O or C-N vibrations, possibly related to the LF-Fe or LF-Alg interaction (Hofstetter et al., 2011). The new band observed at 608 cm^−1^ can be attributed to Fe-O vibration (Stoia et al., 2016), indicating Fe coordination within the system, as it was only present in Fe-containing samples.

No new bands were observed in the FTIR spectra, suggesting the absence of new covalent bond formation. The SDS-PAGE (including reducing conditions) and HPLC analysis however showed that LF and Alg could not be clearly resolved individually in ternary systems (**Fig. S9**). This suggests very strong interactions between LF and Alg which are likely being driven by non-covalent forces such as electrostatic interactions and hydrogen bonding.

#### 3.2.5 Microstructure

SEM images of the freeze-dried samples revealed the microstructure and morphology of the LAF complexes and hydrogels (**Fig. 6)**. Representative samples (LF:Alg - 8:2, 5:5, and 2:8 at both 20 mM and 30 mM Fe(II)) were selected to reflect different structural states encountered.

**Fig. 6.**
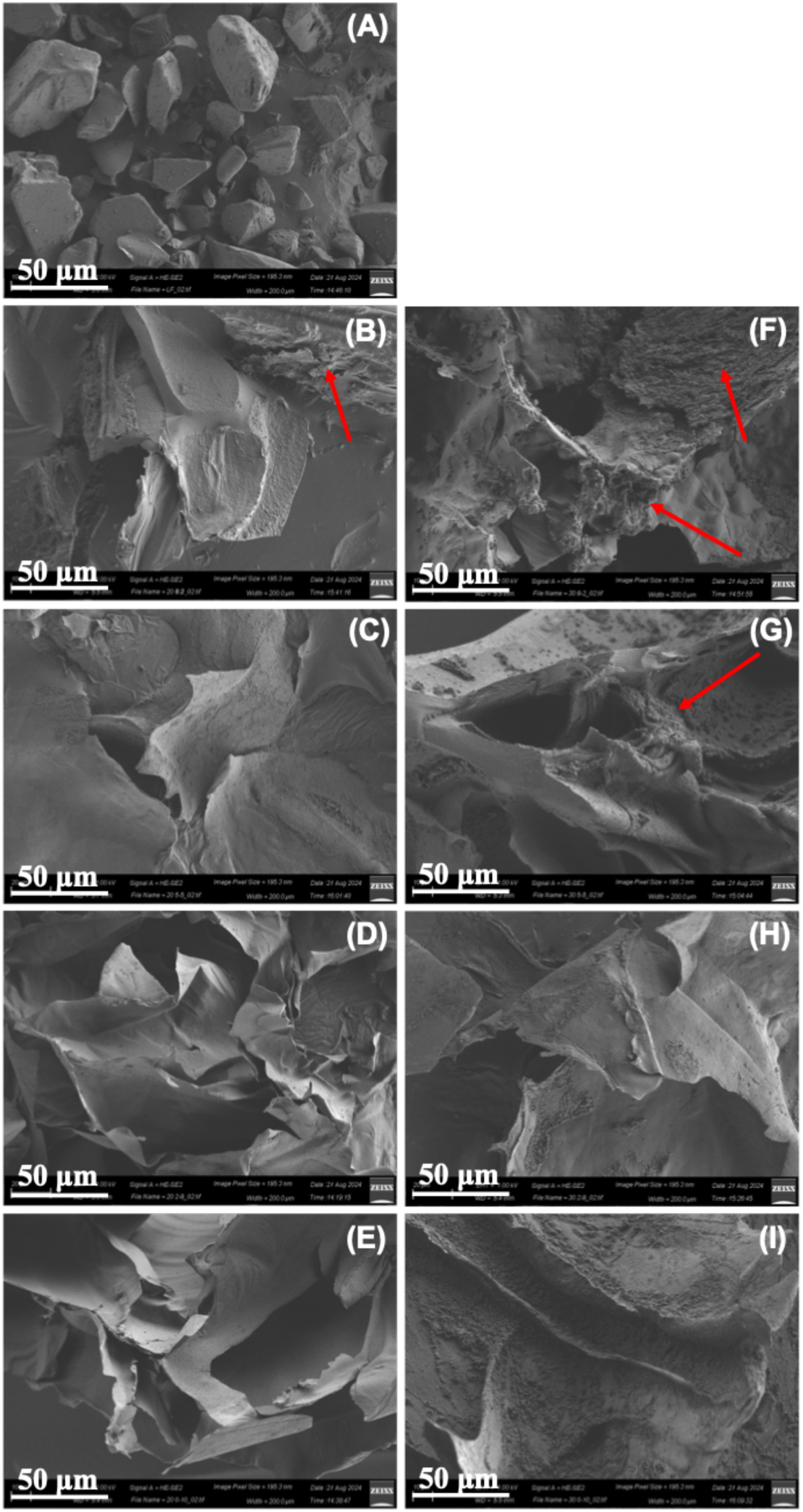
SEM images of LF powder and freeze-dried LAF external surfaces. Scale bar = 50 µm. Red arrows denote electrostatic complexation. (**A**: LF; **B-E:** LAF 8:2, 5:5, 2:8, 0:10 20 mM; **F-I:** LAF 8:2, 5:5, 2:8, 0:10 30 mM)

LF powder appeared as dispersed particles with smooth surface morphology (**Fig. 6A**). The freeze-dried Alg and Alg-Fe 10 mM showed film structure, lacking notable 3D-structure (**Fig. S10A-B**) . With the incorporation of both LF and additional Fe, the morphology of the ternary system showed distinct morphologies depending on the LF/Alg ratio. At higher LF ratios, the system was dominated by complexes, leading to irregular aggregates, as highlighted by red arrows in selected images (**Fig. 6B&F-G**). This observation is consistent with LF-Alg complexes prepared without Fe at lower concentration. Vacuum dried electrostatic complexation samples showed separate particles, and freeze-dried complexes formed bulky aggregates with a rough surface (**Fig. S10C-D**). This suggests the absence of a continuous network structure without Fe-mediated cross-linking. In LAF complexes with LF to Alg ratios of 8:2 and 7:2, aggregates and some thin, sheet-like network structures were observed, suggesting the onset of hydrogel network formation (**Fig. 6B&F-G**). As the proportion of Alg increased, these irregular aggregates gradually diminished and were replaced by more layered, mesh-like structures, consistent with the formation of the hydrogel network.

At a lower Fe(II) concentration (20 mM), the LF to Alg ratio of 8:2 formed a more developed network structure (**Fig. 6B**), whereas at a higher Fe(II) concentration (30 mM), the same composition exhibited more notable aggregate features (**Fig. 6F**). This observation adds to the evidence suggesting that the balance between complexation and Fe(II)-mediated cross-linking is sensitive to both LF to Alg ratio and Fe(II) concentration.

### 3.3 Thermal stability of LAF complexes and hydrogels

Following the formation and characterization of LAF complexes and hydrogels, their thermal stability was evaluated. Differential scanning calorimetry (DSC) was used to assess protein denaturation behavior in the freeze-dried state, while circular dichroism (CD) was used to assess secondary structure changes in rehydrated samples before and after heating. In addition, Fe(II) oxidation behavior was analyzed before and after sample heating.

#### 3.3.1 DSC analysis of LAF powders

The DSC thermograms of LF and LAF complexes and hydrogels showed endothermic peaks typically correspond to protein unfolding and denaturation (**Fig. S11 and Table S2)**, where T_onset_ indicates the onset of denaturation, and T_m_ represents the temperature of maximum heat absorption (Gill et al., 2010). Only samples with well-defined and reproducible thermal transitions are presented. These include native LF and selected LAF formulations (LF:Alg = 8:2, 5:5, and 2:8 at 30 mM Fe(II), and 5:5 at 20 mM Fe(II)).

For LF, two peaks were observed at approximately 61.2 °C and 85.5 °C, corresponding to the N- and C-lobes of the protein structure, respectively, consistent with denaturation (Lin et al., 2022b). The relatively low first T_m_ (∼60 °C) of native LF indicates limited thermal stability, which restricts its applications in food processing.

In LAF composites, the first denaturation peak shifted significantly to higher temperatures (77–82 °C), indicating improved thermal stability. This enhancement can be attributed to interactions between LF and Alg, including electrostatic interactions and hydrogen bonding. LF to Alg ratios of 2:8 and 8:2 exhibited higher T_m_ values, suggesting stronger interactions and improved structural stability. In contrast, LF to Alg ratios 5:5 showed comparatively lower T_m_ values (< 80 °C), consistent with weaker interactions in this composition.

This trend was further supported by the observed enthalpy change (ΔH), where LF to Alg ratios of 2:8 and 8:2 exhibited slightly higher ΔH values than LF alone, indicating increased energy required for unfolding. In contrast, the 5:5 formulations showed comparable or slightly lower ΔH values, and no clear difference was observed between the 20 and 30 mM Fe(II) samples at this ratio. These results suggest enhanced thermal stability of LF after composites formation; however, the DSC data alone do not distinguish the relative contributions of LF–Alg interactions and Fe-mediated cross-linking.

The second denaturation temperature of LF showed no significant endothermic peak within the measurement range, possibly due to its subtle nature. Additionally, as the Alg ratio increased, a noticeable glass transition peak emerged around 133 °C (indicated by dashed arrows in **Fig. S11**), which aligns with previous studies (Espíndola et al., 2023).

#### 3.3.2 Protein secondary structure change of rehydrated LAF after thermal treatment

Rehydrated LAF samples were heated at 95 °C for 5 min at pH 7.0, and structural changes were analyzed by CD spectroscopy (**Fig. 7)**.

**Fig. 7.**
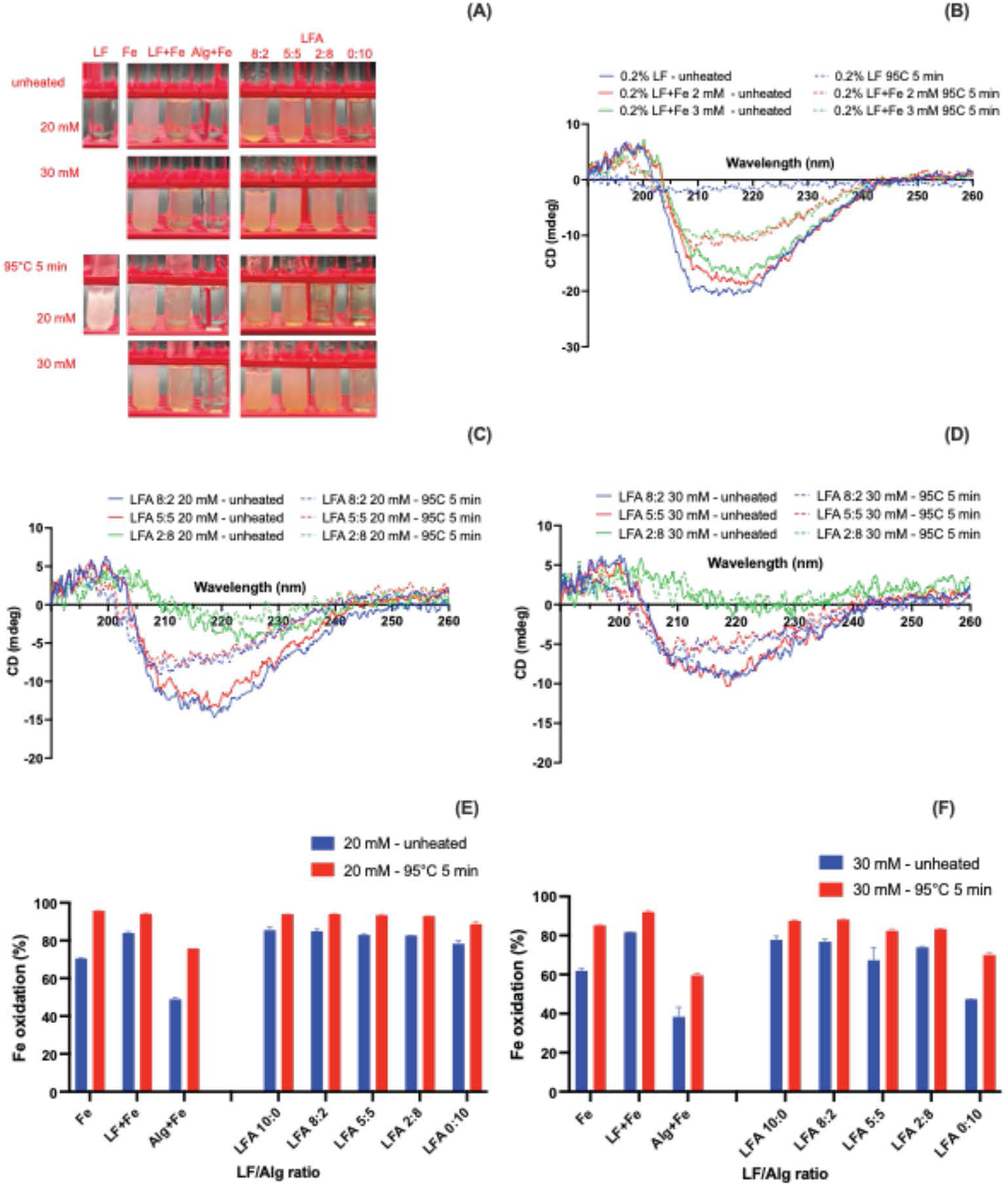
The image of protein and LFA hydrogels before and after heating (A) and their CD spectra at different LF-Alg ratios and Fe concentrations (B: LF and LF+Fe(II) control: C: Fe 20 mM; C: Fe 30 mM).

LF was thermally unstable and underwent noticeable aggregation after heating (**Fig. 7A**), showing almost complete loss of CD signal and indicating extensive unfolding and disruption of secondary structure (**Fig. 7B**). Before heating, LF consists of approximately 50% α-helix and ∼30% β-sheets, whereas after thermal treatment, the α-helix content was nearly completely lost indicating a transition to a disordered structure and protein denaturation (**Fig. S12**). To distinguish the effect of Fe(II) alone from that of the ternary system, 0.2 w/w% LF with Fe(II) (2 and 3 mM) was also examined. The addition of Fe(II) alone resulted in slight decrease in CD signal intensity before heating and provided limited protection after heating, as substantial signal loss was still observed.

In contrast, LAF complexes and hydrogels both showed no severe aggregation after heating, and the CD spectra retained the characteristic features of intact LF, particularly the positive peak around 190 nm associated with α-helical structure (**Fig. 7C-D**).

The extent of structural preservation depended on system composition. At LF to Alg ratios of 8:2 and 5:5, α-helix content showed minimal change after heating, while some β-sheet structures transitioned into disordered conformations (**Fig. S12**), suggesting partial but not complete structural disruption. At higher Fe(II) concentrations, a slight increase in disordered structures was observed, indicating that excess Fe(II) may induce limited conformational rearrangements by binding with the carboxyl or hydroxyl group of protein (Ding et al., 2024).

At the LF:Alg ratio of 2:8, a markedly different structural profile was observed. The α-helix content was nearly absent, while β-sheet structures dominated (>50%), accompanied by a substantial proportion of disordered conformations (∼20%, **Fig. 7C-D**). This structural profile remained largely unchanged after heating, indicating that the system had undergone substantial structural reorganization prior to thermal treatment but was stable during heating.

In contrast, LF with Fe(II) alone showed only minor variations in CD spectra before heating but exhibited noticeable helix loss to disordered structure after heating (**Fig. 7B & S12**), indicating that Fe(II) binding alone does not effectively prevent thermal denaturation. This also suggests that the pronounced structural reorganization at the 2:8 ratio cannot be attributed to Fe(II) binding alone. Instead, it is likely associated with the combined effects of the Alg-rich environment and Fe-mediated network formation, which may impose spatial constraints on LF and promote structural rearrangement toward β-sheet and disordered conformations.

Despite these changes, LAF complexes and hydrogels maintained greater structural integrity compared to free LF or LF-Fe(II) mixtures, demonstrating effective protection against thermal denaturation. This enhanced stability is likely attributed to the combined effects of LF-Alg interactions and Alg-Fe cross-linked networks, which physically confine LF within the matrix and restrict molecular mobility, thereby suppressing unfolding and aggregation during thermal treatment.

#### 3.3.3 Fe(II) oxidation of rehydrated LAF complexes and hydrogels before and after thermal treatment

Free Fe(II) showed substantial oxidation and reduced solubility at neutral pH, with over 60% oxidized before heating and up to 80-90% after heating (**Fig. 7A, E&F**).

In the presence of either LF or Alg, Fe(II) precipitation was reduced, indicating improved dispersion. This can be attributed to Fe(II) coordination with LF and Alg. However, LF alone did not effectively prevent Fe(II) oxidation, while alginate provided partial protection, consistent with previous reports (Katuwavila et al., 2016).

At neutral conditions, over 80% of the Fe(II) was already oxidized before heating in rehydrated LAF complexes and hydrogels at 20 mM Fe(II), much higher than in samples dissolved at acidic conditions (<20%, **Fig. 5)**. As the Fe(II) concentration increased, particularly in samples with higher Alg ratios, the percentage of Fe(II) oxidation decreased. Specifically, in the LF to Alg ratio of 0:10 and a Fe(II) concentration of 30 mM at neutral pH, about 40% of the Fe(II) was oxidized before heating, increasing to 70% after heating. This can be attributed to the stronger cross-linking between Fe and Alg, in which the Fe(II) was better encapsulated within the egg-box structure, reducing its exposure to oxygen and thus limiting oxidation. When compared with the mixture of Alg and Fe(II), the oxidation rate of LF to Alg ratio 0:10 was higher than the direct mixture of Alg and Fe(II). This might be due to the freeze-dry process, which has been reported to enhance the oxidation of the bioactive ingredients in the food matrix, as the higher porosity and lower water content of freeze dried samples make them more accessible to oxygen (Silva-Espinoza et al., 2019).

Overall, neither LAF complexes nor hydrogels do not effectively protect Fe(II) from oxidation in neutral environments even before heating. Even though the Fe(II) was encapsulated within the Alg blocks in hydrogels, further protection was still needed for Fe(II) oxidation prevention.

## 4 Conclusion

A simple and effective LF–Fe co-encapsulation system was developed through the combination of electrostatic interactions between LF and alginate and Fe-induced alginate gelation. By tuning the LF to Alg ratio and the Fe(II) concentration, the ternary system exhibited distinct structural transitions, ranging from complex at higher LF ratios to hydrogel networks at higher Alg content.

At higher LF ratios, the system was dominated by electrostatic complexation, while increasing Alg content and Fe(II) concentrations promoted the formation of egg-box structures and the development of a more robust three-dimensional hydrogel network. These structures were further stabilized by hydrogen bonding along the Alg chains.

Rheological and swelling analyses confirmed that the stability of the ternary system was significantly enhanced compared to Alg-Fe gels alone, highlighting the critical role of the LF-Alg electrostatic interactions. The system showed composition-dependent behavior, where stronger electrostatic interactions favored LF-rich compositions and Alg-rich compositions favored hydrogen bonding and cross-linking. The combination of these interactions led to improved mechanical strength and stability.

The formation of three-dimensional network structures in Alg-rich systems and aggregated complex structures at higher LF ratios were observed in the SEM and FTIR analyses indicating that the interactions within the system were primarily non-covalent, allowing the polymers to retain their structural integrity.

Thermal studies further demonstrated that the LAF system effectively protected LF against thermal denaturation, preserving secondary structure compared to LF alone. However, the system showed limited ability to prevent Fe(II) oxidation under neutral conditions, highlighting the dominant influence of pH on iron stability.

Overall, the LAF ternary system enable high Fe(II) loading, improves protein thermal stability, and provides a promising platform for co-delivery applications. With further optimization of iron stability and release behavior, this system has potential for use in iron-fortified and high-protein functional food applications.

## CRediT authorship contribution statement

**Yunan Huang**: Conceptualization, Investigation, Data curation, Formal analysis, Writing – original draft, Writing – review & editing. **Tiantian Lin**: Conceptualization, Methodology, Investigation, Writing – review & editing. **Waritsara Khongkomolsakul**: Investigation, Writing – review & editing. **Jieying Li**: Formal analysis, Writing – review & editing. **Claire Elizabeth Noack**: Investigation, Writing – review & editing. **Younas Dadmohammadi**: Supervision, Funding acquisition, Writing – review & editing. **Alireza Abbaspourrad**: Project Administration, Funding acquisition, Resources, Writing – review & editing, Supervision.

## Declaration of Competing Interest

All authors declare no conflict of interests to influence the work reported in this paper.

## Generative AI Statement

The authors certify that generative AI was not used in preparing this article. Non-generative AI, such as spelling and grammar checkers in Office 365 and Google Docs, and citation management software, was used. All instances when non-generative AI was used were reviewed by the authors and editors.

## Acknowledgements

This project was funded by the Bill & Melinda Gates Foundation (INV-039533). The authors acknowledge the use of facilities and instrumentation supported by NSF through the Cornell University Materials Research Science and Engineering Center DMR-1719875.

## Appendix A. Supplementary data

The following are the Supplementary data to this article:

## Data availability

## Notes

https://doi.org/10.5281/zenodo.19683185

### Competing Interest Statement

The authors have declared no competing interest.

## Reference

Barth, A., & Zscherp, C. (2002). What vibrations tell us about proteins. Quarterly Reviews of Biophysics, 35(4), 369–430. 10.1017/s0033583502003815

Bastos, L. P. H., De Carvalho, C. W. P., & Garcia-Rojas, E. E. (2018). Formation and characterization of the complex coacervates obtained between lactoferrin and sodium alginate. International Journal of Biological Macromolecules, 120, 332–338. 10.1016/j.ijbiomac.2018.08.050

Bengoechea, C., Peinado, I., & McClements, D. J. (2011). Formation of protein nanoparticles by controlled heat treatment of lactoferrin: Factors affecting particle characteristics. Food Hydrocolloids, 25(5), 1354–1360. 10.1016/j.foodhyd.2010.12.014

Bokkhim, H., Bansal, N., GrØndahl, L., & Bhandari, B. (2013). Physico-chemical properties of different forms of bovine lactoferrin. Food Chemistry, 141(3), 3007–3013. 10.1016/j.foodchem.2013.05.139

Bokkhim, H., Bansal, N., Grøndahl, L., & Bhandari, B. (2015). Interactions between different forms of bovine lactoferrin and sodium alginate affect the properties of their mixtures. Food Hydrocolloids, 48, 38–46. 10.1016/j.foodhyd.2014.12.036

Breymann, C. (2015). Iron Deficiency Anemia in Pregnancy. *Seminars in Hematology*, Anemia in Clinical Practice, 52(4), 339–347. 10.1053/j.seminhematol.2015.07.003

Cofelice, M., Messia, M. C., Marconi, E., Cuomo, F., & Lopez, F. (2023). Effect of the xanthan gum on the rheological properties of alginate hydrogels. Food Hydrocolloids, 142, 108768. 10.1016/j.foodhyd.2023.108768

Dash, S., Gutti, P., Behera, B., & Mishra, D. (2023). Emergence of Anionic Counterparts of Divalent Metal Salts as the Fine-Tuners of Alginate Hydrogel Properties for Tissue Engineering and Drug Delivery Applications. 10.26434/chemrxiv-2023-rzn2r

Ding, X., Liu, Y., Zheng, L., Chang, Q., Chen, X., & Xi, C. (2024). Effect of different iron ratios on interaction and thermodynamic stability of bound whey protein isolate. Food Research International, 182, 114198. 10.1016/j.foodres.2024.114198

Dyrda-Terniuk, T., & Pomastowski, P. (2023). The Multifaceted Roles of Bovine Lactoferrin: Molecular Structure, Isolation Methods, Analytical Characteristics, and Biological Properties. Journal of Agricultural and Food Chemistry, 71(51), 20500–20531. 10.1021/acs.jafc.3c06887

Espíndola, S. P., Norder, B., Koper, G. J. M., & Picken, S. J. (2023). The Glass Transition Temperature of Heterogeneous Biopolymer Systems. Biomacromolecules, 24(4), 1627–1637. 10.1021/acs.biomac.2c01356

Gholam Jamshidi, E., Behzad, F., Adabi, M., & Esnaashari, S. S. (2024). Edible Iron-Pectin Nanoparticles: Preparation, Physicochemical Characterization and Release Study. Food and Bioprocess Technology, 17(3), 628–639. 10.1007/s11947-023-03156-4

Gill, P., Moghadam, T. T., & Ranjbar, B. (2010). Differential Scanning Calorimetry Techniques: Applications in Biology and Nanoscience. Journal of Biomolecular Techniques : JBT, 21(4), 167–193.

Hofstetter, T. E., Howder, C., Berden, G., Oomens, J., & Armentrout, P. B. (2011). Structural Elucidation of Biological and Toxicological Complexes: Investigation of Monomeric and Dimeric Complexes of Histidine with Multiply Charged Transition Metal (Zn and Cd) Cations using IR Action Spectroscopy. The Journal of Physical Chemistry B, 115(43), 12648–12661. 10.1021/jp207294b

Hong, R., Xie, A., Jiang, C., Guo, Y., Zhang, Y., Chen, J., Shen, X., Li, M., & Yue, X. (2024). A review of the biological activities of lactoferrin: Mechanisms and potential applications. Food & Function, 15(16), 8182–8199. 10.1039/D4FO02083A

Hu, C., Lu, W., Mata, A., Nishinari, K., & Fang, Y. (2021). Ions-induced gelation of alginate: Mechanisms and applications. International Journal of Biological Macromolecules, 177, 578–588. 10.1016/j.ijbiomac.2021.02.086

Huang, Y., Lin, T., Dadmohammadi, Y., He, Y., Khongkomolsakul, W., Noack, C. E., & Abbaspourrad, A. (2024). Lactoferrin thermal stabilization and iron(II) fortification through ternary complex fabrication with succinylated sodium caseinate. Food Chemistry: X, 22, 101498. 10.1016/j.fochx.2024.101498

Hurrell, R. F. (2002). Fortification: Overcoming Technical and Practical Barriers. The Journal of Nutrition, 132(4), 806S–812S. 10.1093/jn/132.4.806S

Hurrell, R. F. (2022). Ensuring the Efficacious Iron Fortification of Foods: A Tale of Two Barriers. Nutrients, 14(8). 10.3390/nu14081609

Jenssen, H., & Hancock, R. E. W. (2009). Antimicrobial properties of lactoferrin. *Biochimie*, Advances in Lactoferrin Research, 91(1), 19–29. 10.1016/j.biochi.2008.05.015

Katuwavila, N. P., Perera, A. D. L. C., Dahanayake, D., Karunaratne, V., Amaratunga, G. A. J., & Karunaratne, D. N. (2016). Alginate nanoparticles protect ferrous from oxidation: Potential iron delivery system. International Journal of Pharmaceutics, 513(1), 404–409. 10.1016/j.ijpharm.2016.09.053

Kumari, A., & Chauhan, A. K. (2022). Iron nanoparticles as a promising compound for food fortification in iron deficiency anemia: A review. Journal of Food Science and Technology, 59(9), 3319–3335. 10.1007/s13197-021-05184-4

Li, J., & Mooney, D. J. (2016). Designing hydrogels for controlled drug delivery. Nature Reviews Materials, 1(12), Article 12. 10.1038/natrevmats.2016.71

Lin, T., Dadmohammadi, Y., Davachi, S. M., Torabi, H., Li, P., Pomon, B., Meletharayil, G., Kapoor, R., & Abbaspourrad, A. (2022a). Improvement of lactoferrin thermal stability by complex coacervation using soy soluble polysaccharides. Food Hydrocolloids, 131, 107736. 10.1016/j.foodhyd.2022.107736

Lin, T., Dadmohammadi, Y., Davachi, S. M., Torabi, H., Li, P., Pomon, B., Meletharayil, G., Kapoor, R., & Abbaspourrad, A. (2022b). Improvement of lactoferrin thermal stability by complex coacervation using soy soluble polysaccharides. Food Hydrocolloids, 131, 107736. 10.1016/j.foodhyd.2022.107736

Lin, T., & Fernández-Fraguas, C. (2020). Effect of thermal and high-pressure processing on the thermo-rheological and functional properties of common bean (*Phaseolus vulgaris* L.) flours. LWT, 127, 109325. 10.1016/j.lwt.2020.109325

Lin, T., Zhou, Y., Dadmohammadi, Y., Yaghoobi, M., Meletharayil, G., Kapoor, R., & Abbaspourrad, A. (2023). Encapsulation and stabilization of lactoferrin in polyelectrolyte ternary complexes. Food Hydrocolloids, 145, 109064. 10.1016/j.foodhyd.2023.109064

Liu, F., Zhang, S., Li, J., McClements, D. J., & Liu, X. (2018). Recent development of lactoferrin-based vehicles for the delivery of bioactive compounds: Complexes, emulsions, and nanoparticles. Trends in Food Science & Technology, 79, 67–77. 10.1016/j.tifs.2018.06.013

McAvan, B. S., France, A. P., Bellina, B., Barran, P. E., Goodacre, R., & Doig, A. J. (2020). Quantification of protein glycation using vibrational spectroscopy. Analyst, 145(10), 3686–3696. 10.1039/C9AN02318F

Mikulic, N., Uyoga, M. A., Mwasi, E., Stoffel, N. U., Zeder, C., Karanja, S., & Zimmermann, M. B. (2020). Iron Absorption is Greater from Apo-Lactoferrin and is Similar Between Holo-Lactoferrin and Ferrous Sulfate: Stable Iron Isotope Studies in Kenyan Infants. The Journal of Nutrition, 150(12), 3200–3207. 10.1093/jn/nxaa226

Miles, A. J., Ramalli, S. G., & Wallace, B. A. (2022). DichroWeb, a website for calculating protein secondary structure from circular dichroism spectroscopic data. Protein Science, 31(1), 37–46. 10.1002/pro.4153

Morgan, B., & Lahav, O. (2007). The effect of pH on the kinetics of spontaneous Fe(II) oxidation by O2 in aqueous solution – basic principles and a simple heuristic description. Chemosphere, 68(11), 2080–2084. 10.1016/j.chemosphere.2007.02.015

Noack, C. E., Lin, T., Dadmohammadi, Y., Huang, Y., & Abbaspourrad, A. (2025). Improved thermal stability of lactoferrin and oxidative stability of Iron(II) sulfate by Co-encapsulation with low-methoxyl pectin. Food Hydrocolloids, 169, 111570. 10.1016/j.foodhyd.2025.111570

Rao, M. A. A. (2010). Rheology of Fluid and Semisolid Foods: Principles and Applications. Springer Science & Business Media.

Rosa, L., Cutone, A., Lepanto, M. S., Paesano, R., & Valenti, P. (2017). Lactoferrin: A Natural Glycoprotein Involved in Iron and Inflammatory Homeostasis. International Journal of Molecular Sciences, 18(9), Article 9. 10.3390/ijms18091985

Santiago, P. (2012). Ferrous versus Ferric Oral Iron Formulations for the Treatment of Iron Deficiency: A Clinical Overview. The Scientific World Journal, 2012, e846824. 10.1100/2012/846824

Shen, Y., Posavec, L., Bolisetty, S., Hilty, F. M., Nyström, G., Kohlbrecher, J., Hilbe, M., Rossi, A., Baumgartner, J., Zimmermann, M. B., & Mezzenga, R. (2017). Amyloid fibril systems reduce, stabilize and deliver bioavailable nanosized iron. Nature Nanotechnology, 12(7), Article 7. 10.1038/nnano.2017.58

Silva-Espinoza, M. A., Ayed, C., Foster, T., Camacho, M. del M., & Martínez-Navarrete, N. (2019). The Impact of Freeze-Drying Conditions on the Physico-Chemical Properties and Bioactive Compounds of a Freeze-Dried Orange Puree. Foods, 9(1), 32. 10.3390/foods9010032

Stoia, M., Istratie, R., & Păcurariu, C. (2016). Investigation of magnetite nanoparticles stability in air by thermal analysis and FTIR spectroscopy. Journal of Thermal Analysis and Calorimetry, 125(3), 1185–1198. 10.1007/s10973-016-5393-y

Theurl, I., Aigner, E., Theurl, M., Nairz, M., Seifert, M., Schroll, A., Sonnweber, T., Eberwein, L., Witcher, D. R., Murphy, A. T., Wroblewski, V. J., Wurz, E., Datz, C., & Weiss, G. (2009). Regulation of iron homeostasis in anemia of chronic disease and iron deficiency anemia: Diagnostic and therapeutic implications. Blood, 113(21), 5277–5286. 10.1182/blood-2008-12-195651

