## Supplemental Figures and Tables for "Iron-mediated assembly of lactoferrin-alginate composites for iron encapsulation and structural stabilization"

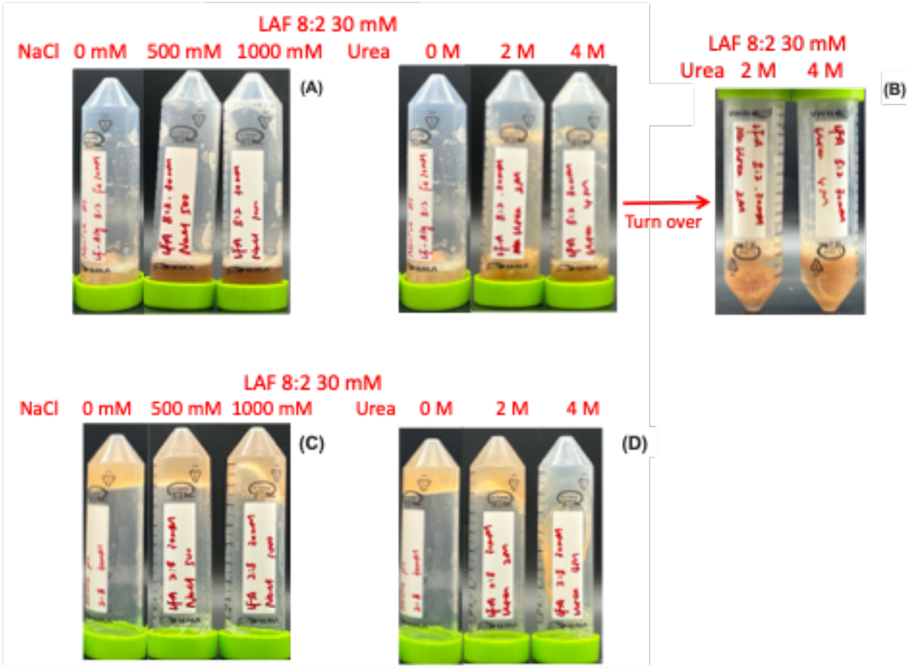


Fig. S1. Hydrogel formation for LAF 8:2 and 2:8 30 mM hydrogel under different blocking agents (A, F: NaCl; B, D: urea).

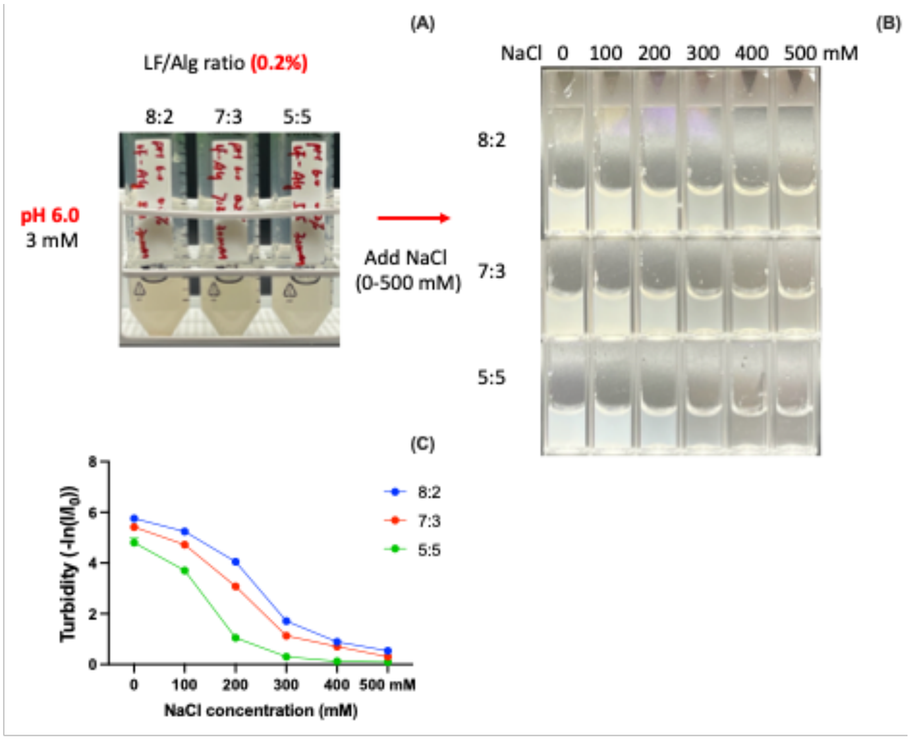


Fig. S2. Electrostatic complex formation of LF-Alg systems at low concentration and its disruption by NaCl. A: complex formation; B: complex mixtures with different NaCl concentration added; C: Turbidity change with NaCl adding.

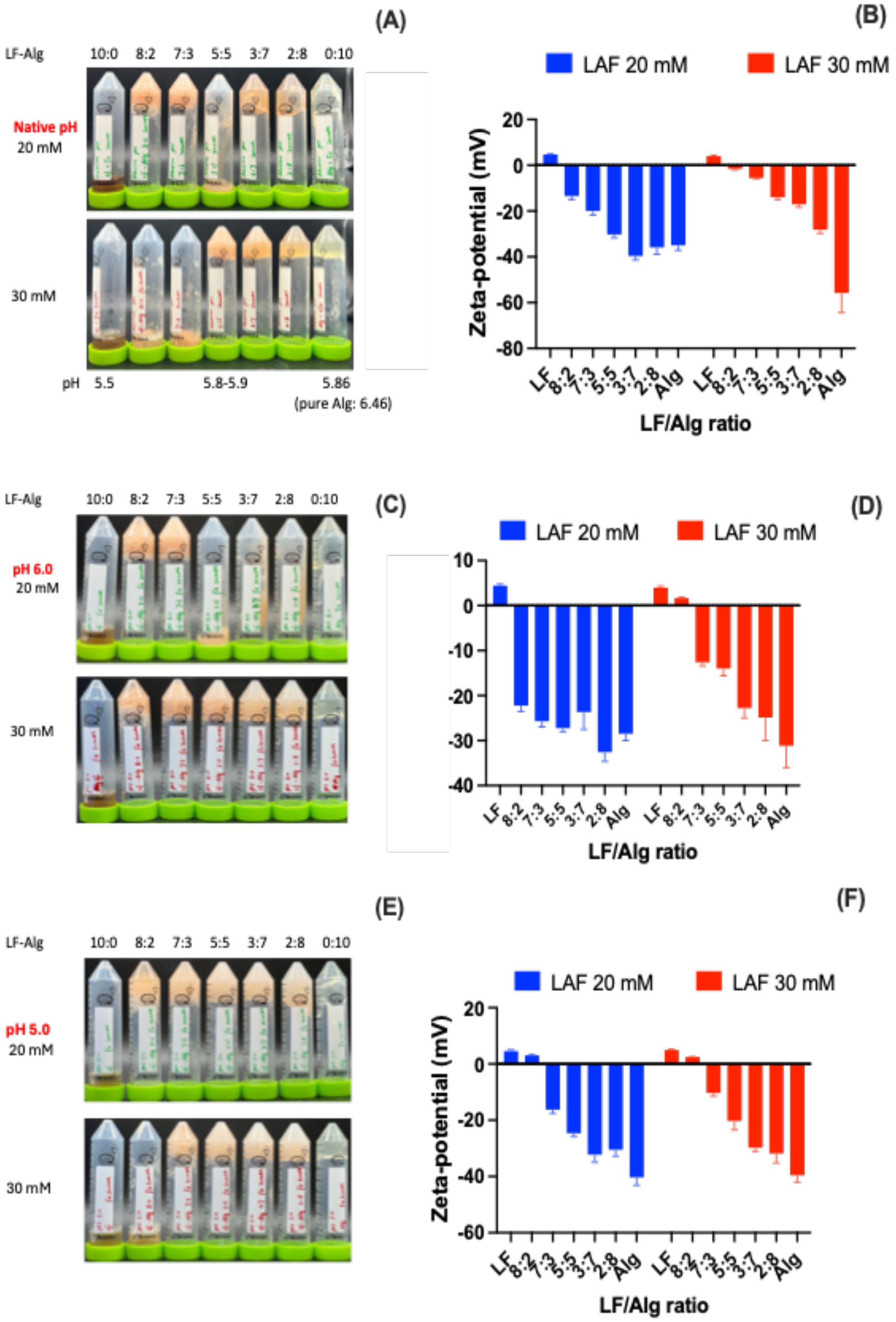


Fig. S3. LFA formation condition at different pHs (A, C, E) and the corresponding zeta-potential (B, D, F) of the hydrogels with different LF-Alg ratios and Fe concentrations (A, B: native pH; C, D: pH 6.0; E, F: pH 5.0).

Table S1: Parameters fitted by Power Law model in oscillation frequency sweep of representative LAF samples.

| | LF/Alg ratio | $lnG'=lnG'_0 + n'ln\omega$ | | | $lnG''=lnG''_0 + n''ln\omega$ | | | $G'_0$-$G''_0$ (Pa) |
|---|---|---|---|---|---|---|---|---|
| | | $lnG'_0$ (Pa) | n' | $R^2$ | $lnG''_0$ (Pa) | n'' | $R^2$ | |
| LAF 20 mM | 8:2 | 3.908 | 0.087 | 0.959 | 3.012 | 0.092 | 0.877 | 7063.764 |
| | 7:3 | 3.648 | 0.115 | 0.970 | 2.943 | 0.021 | 0.882 | 3571.526 |
| | 5:5 | 0.636 | 0.261 | 0.906 | 0.228 | 0.202 | 0.974 | 2.638 |
| | 3:7 | 1.569 | 0.102 | 0.950 | 0.781 | 0.289 | 0.894 | 31.068 |
| | 2:8 | 1.606 | 0.084 | 0.972 | 0.860 | 0.362 | 0.942 | 33.097 |
| | 0:10 | 0.448 | 0.492 | 0.977 | 0.566 | 0.745 | 0.957 | -0.877 |
| LAF 30 mM | 8:2 | 3.879 | 0.101 | 0.984 | 3.123 | 0.029 | 0.722 | 6234.965 |
| | 7:3 | 3.813 | 0.105 | 0.968 | 3.063 | 0.022 | 0.289 | 5344.213 |
| | 5:5 | 2.456 | 0.107 | 0.921 | 1.762 | 1.765 | 0.224 | 227.824 |
| | 3:7 | 2.564 | 0.041 | 0.991 | 1.403 | 0.088 | 0.709 | 341.423 |
| | 2:8 | 2.725 | 0.035 | 0.975 | 1.366 | 0.174 | 0.993 | 507.967 |
| | 0:10 | 2.002 | 0.104 | 0.979 | 1.258 | 0.346 | 0.961 | 82.452 |

G' (Pa): storage moduli; G'' (Pa): loss moduli; ω (rad/s): oscillation frequency; n' and n'' represent the dependence degree of both moduli on oscillation frequency; $G'_0$-$G''_0$ represent gel strength.

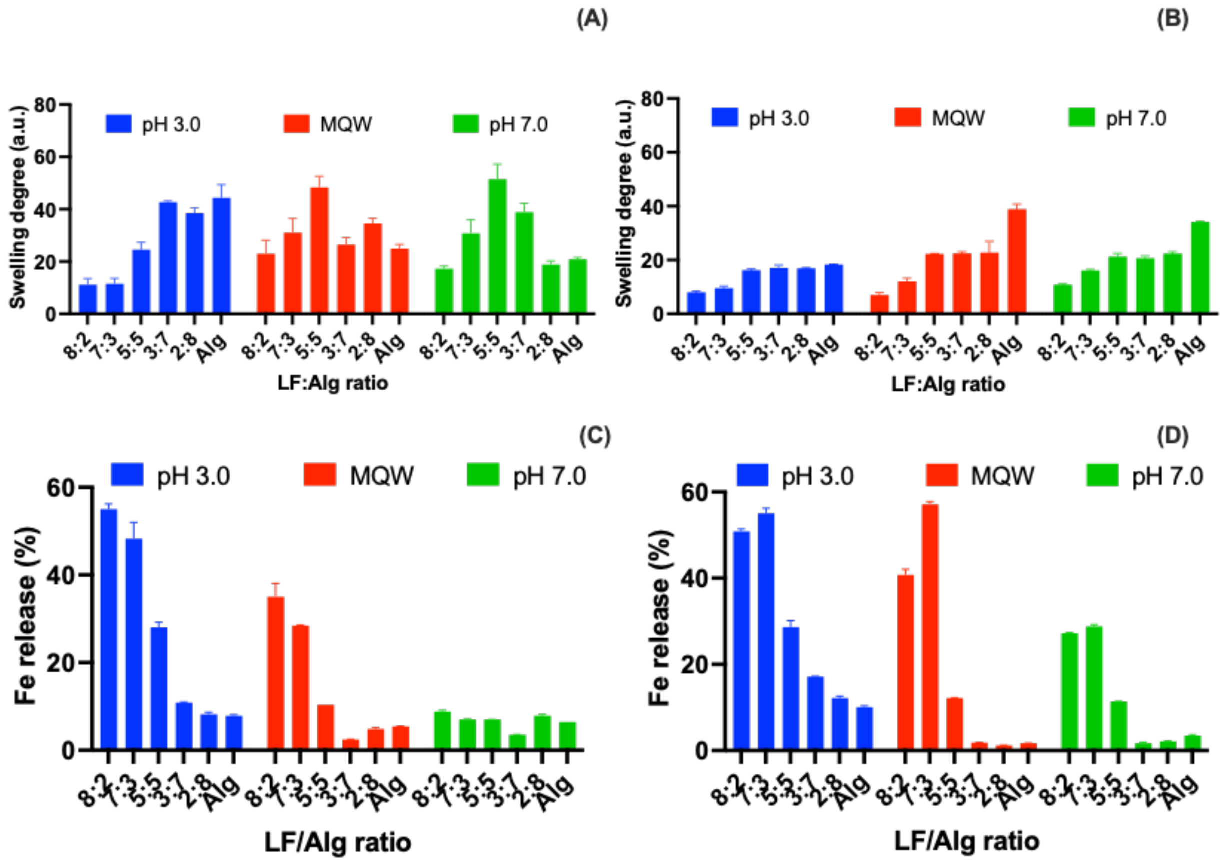


Fig. S4. The swelling degree (A-B) and Fe release (C-D) of LAF with different LF-Alg ratios and Fe concentrations (A,C: 20 mM; B, D:30 mM).

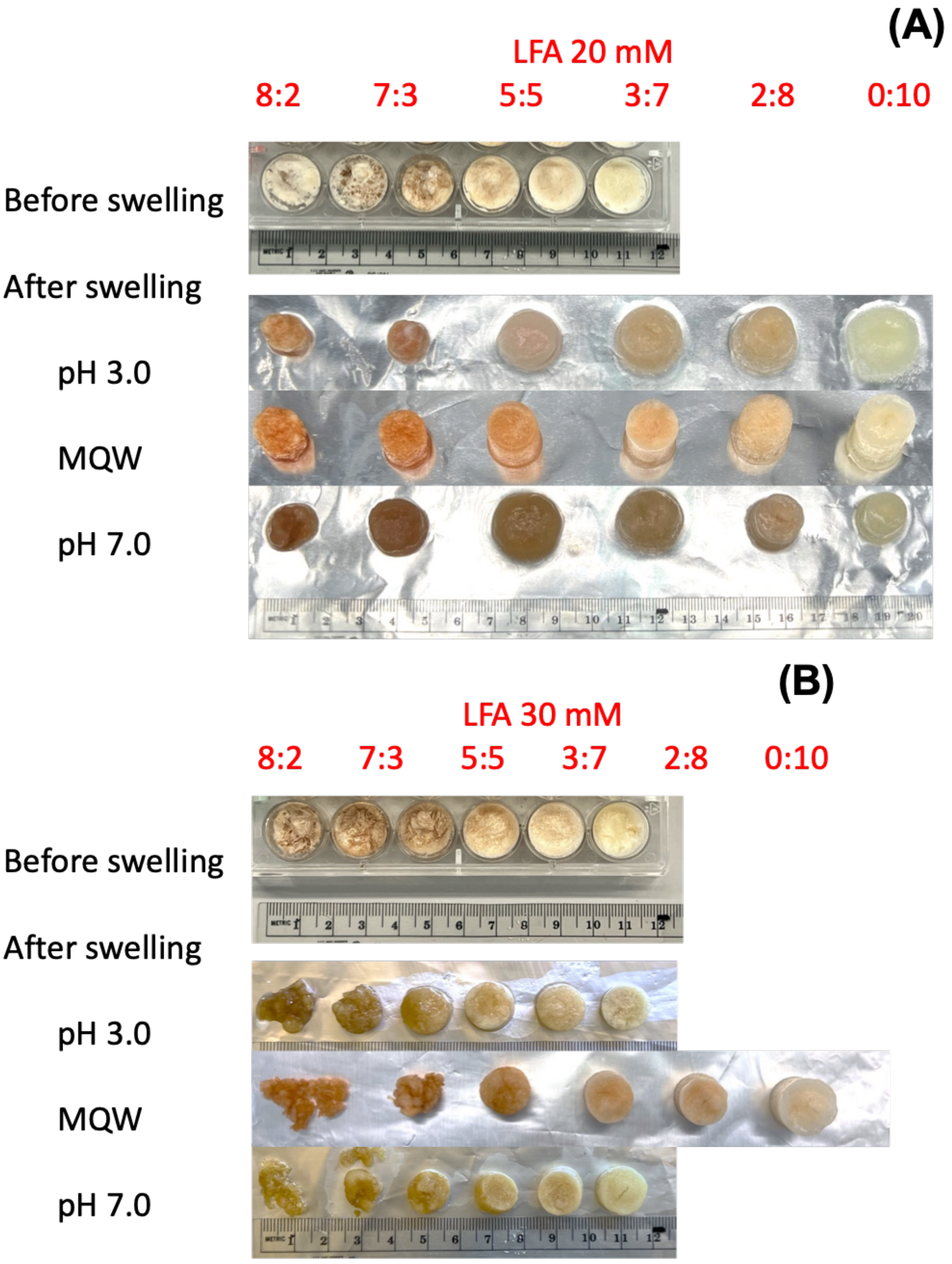


Fig. S5. The image of LAF before and after swelling at different LF-Alg ratios and Fe concentrations in different pHs (A: 20 mM; B: 30 mM). The images are with the same scale and the LF/Alg ratios of each row are from 8:2 to 0:10.

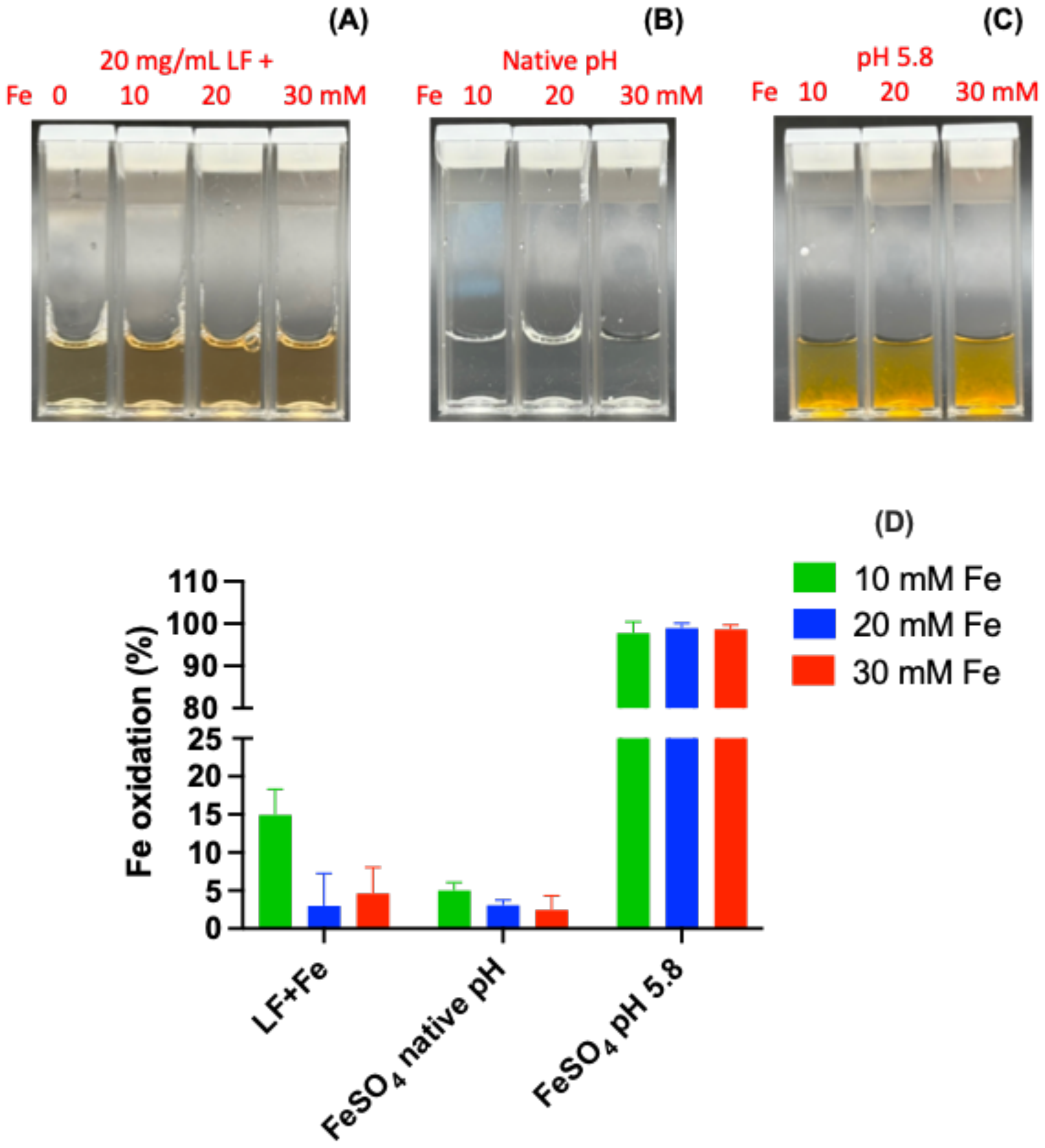


Fig. S6. The image of 20 mg/mL LF + Fe 0, 10, 20 30 mM (A, the pH of LF+Fe mixture ~5.8-5.9), the 10-30 mM $FeSO_4$ solution under native pH (B, pH ~3.2-3.4) and pH 5.8 (C) and the corresponding Fe oxidation (D).

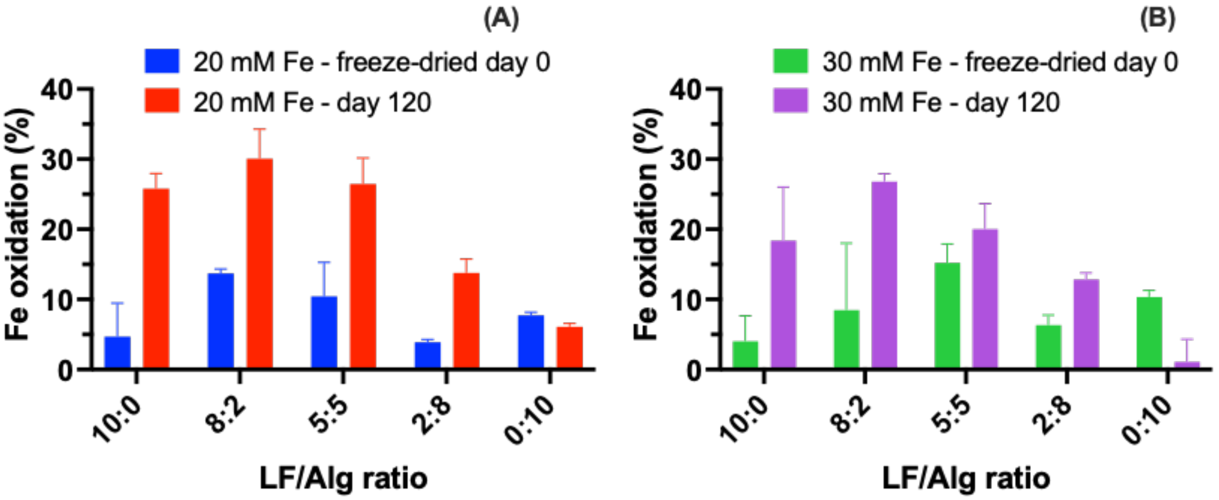


Fig. S7. Storage stability of iron. Fe oxidation of freeze-dried LAF powders on day 0 and day 120 at different LF-Alg ratios and Fe concentrations (A: 20 mM; B; 30 mM).

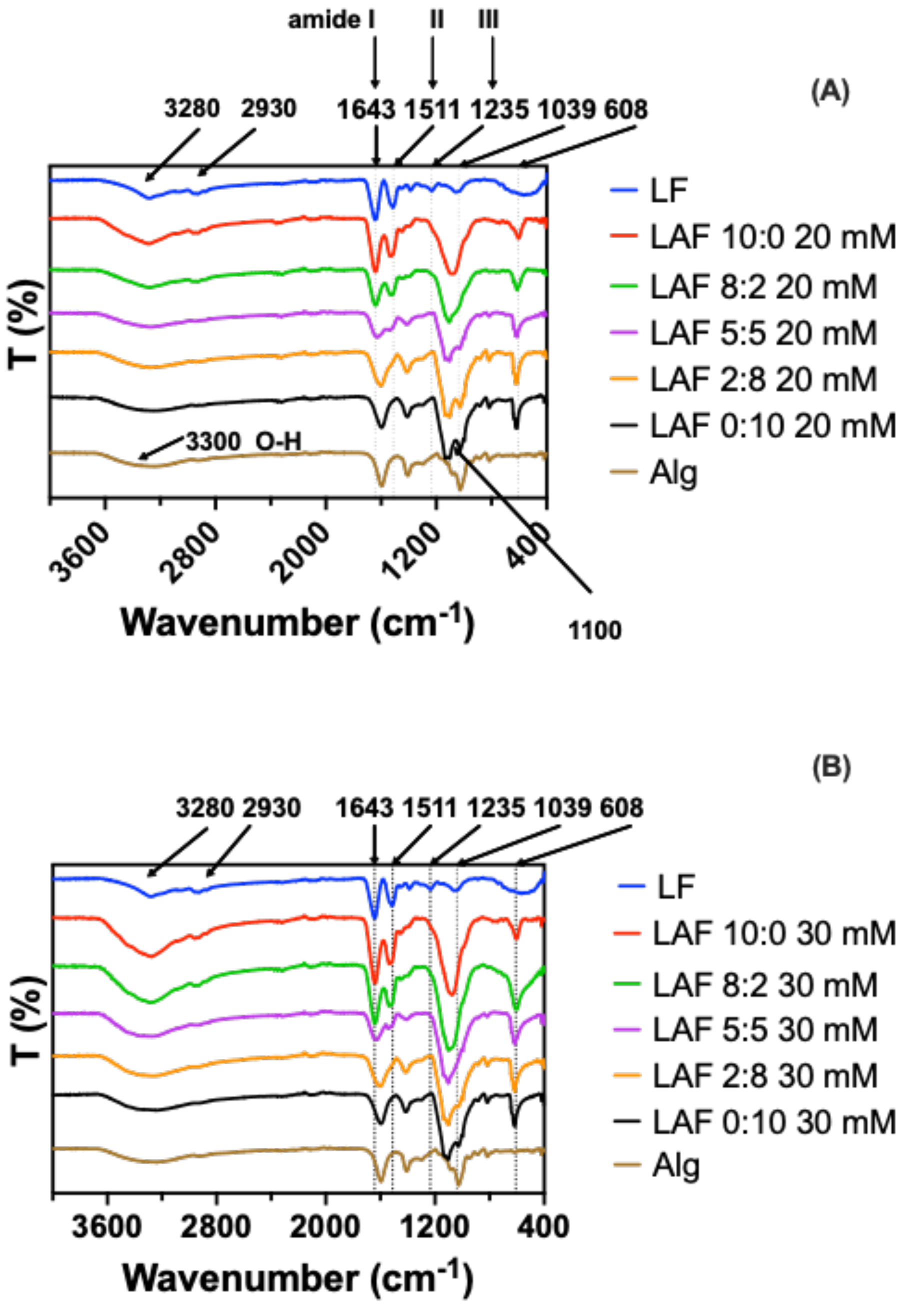


Fig. S8. FT-IR of polymers and the freeze-dried LAF at different LF-Alg ratios and Fe concentrations (A: 20 mM; B: 30 mM).

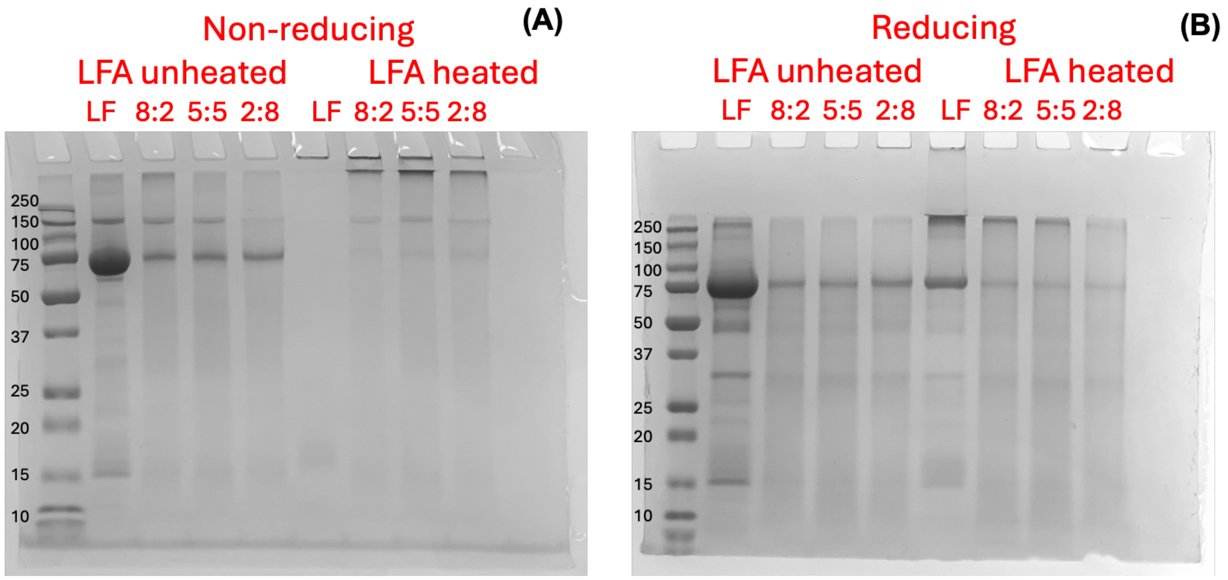


Fig. S9. Non-reducing (A) and reducing (B) SDS-PAGE of LF/Fe mixture and freeze-dried LAF (30 mM Fe(II)) at different LF/Alg ratios. All samples were normalized to the same LF concentration before loading.

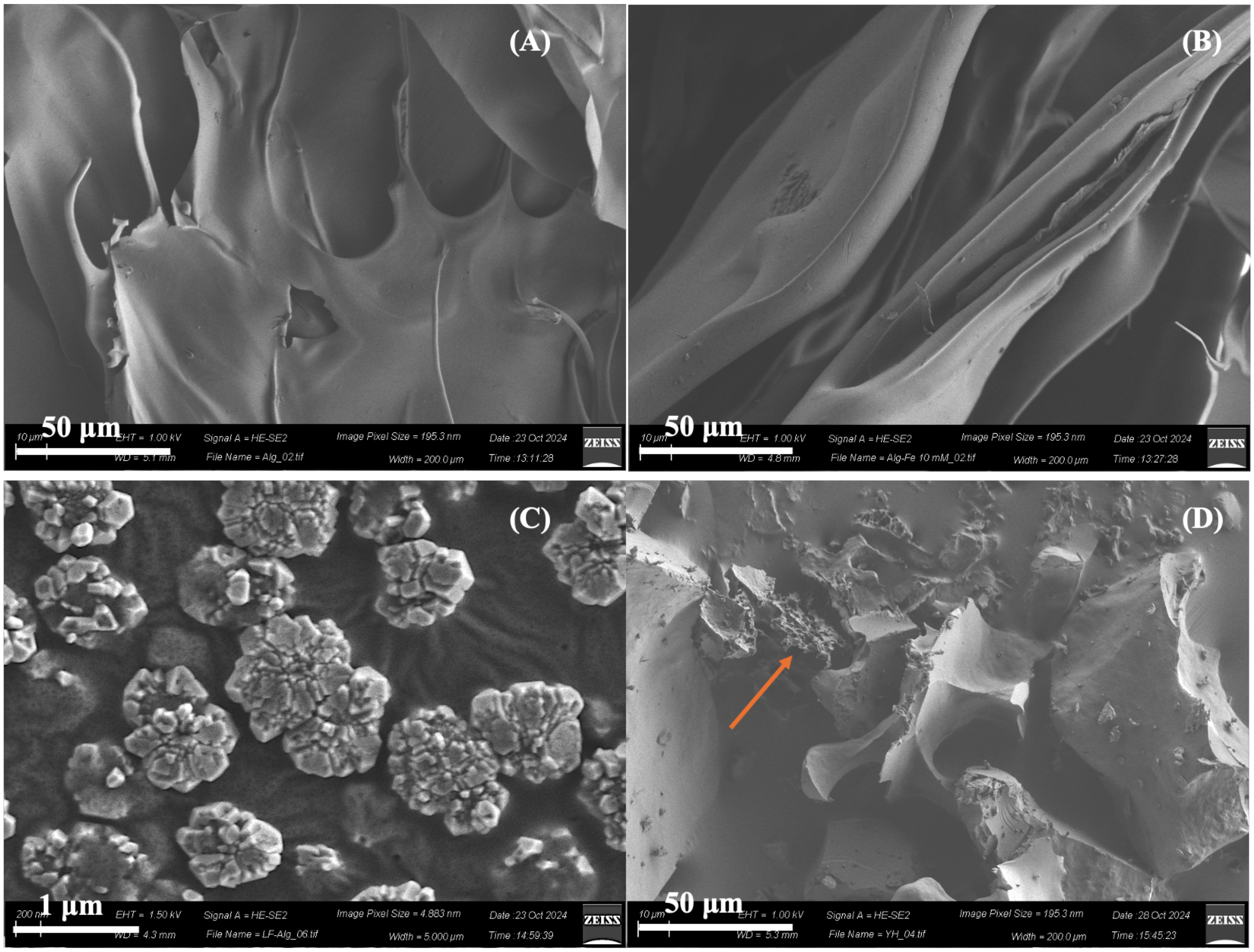


Fig. S10. SEM images of freeze-dried Alg (A), Alg-Fe 10 mM (B), and LF-Alg electrostatic complexation sample. (C): vacuum-dried, scale bar = 1 µm; (D): freeze-dried. Scale bar = 50 µm.

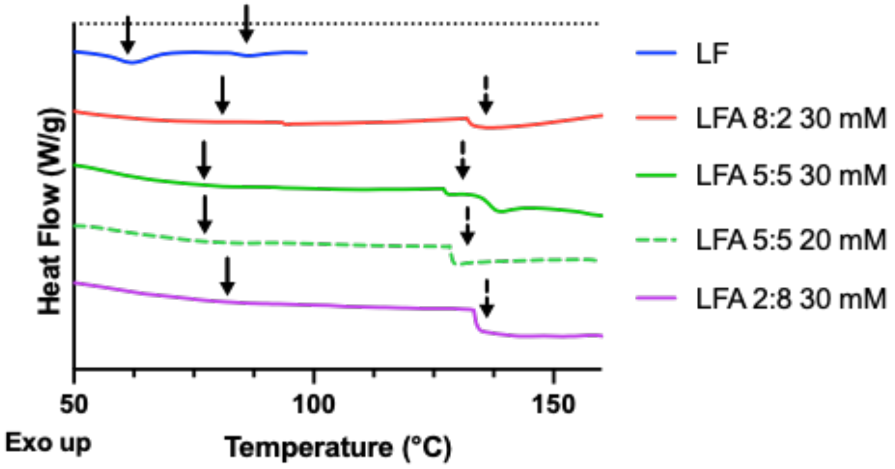


Fig. S11. DSC curve of rehydrated LAF powder at different LF-Alg ratios and Fe concentrations. The arrows indicate the peak temperatures (Tm) and glass transition temperature (Tg) of samples. The curves have been offset for clarity.

Table S2: Thermal properties of LAF samples analyzed from DSC results.

| Sample name | Peak 1 | | | Peak 2 | | |
|---|---|---|---|---|---|---|
| | $T_{onset}$ (°C) | $T_m$ (°C) | Δ*H* (J/g) | $T_{onset}$ (°C) | $T_m$ (°C) | Δ*H* (J/g) |
| Native LF | 52.71 ± 0.76 | 61.57 ± 0.44 | 12.61 ± 0.32 | 81.91 ± 0.30 | 85.49 ± 1.39 | 1.81± 0.0 |
| LAF 8:2 30 mM | 52.27 ± 0.35 | 80.52 ± 0 | 13.45 ± 0.99 | - | - | - |
| LAF 5:5 30 mM | 55.57 ± 0.01 | 76.75 ± 3.70 | 13.62 ± 0.10 | - | - | - |
| LAF 5:5 **20 mM** | 54.81± 0.78 | 77.11 ± 0 | 12.33 ± 2.19 | - | - | - |
| LAF 2:8 30 mM | 51.64 ± 0.68 | 81.71 ± 0 | 14.02 ± 0.68 | - | - | - |

$T_{onset}$ : onset temperature; $T_m$: peak temperature; ΔH : enthalpy change.

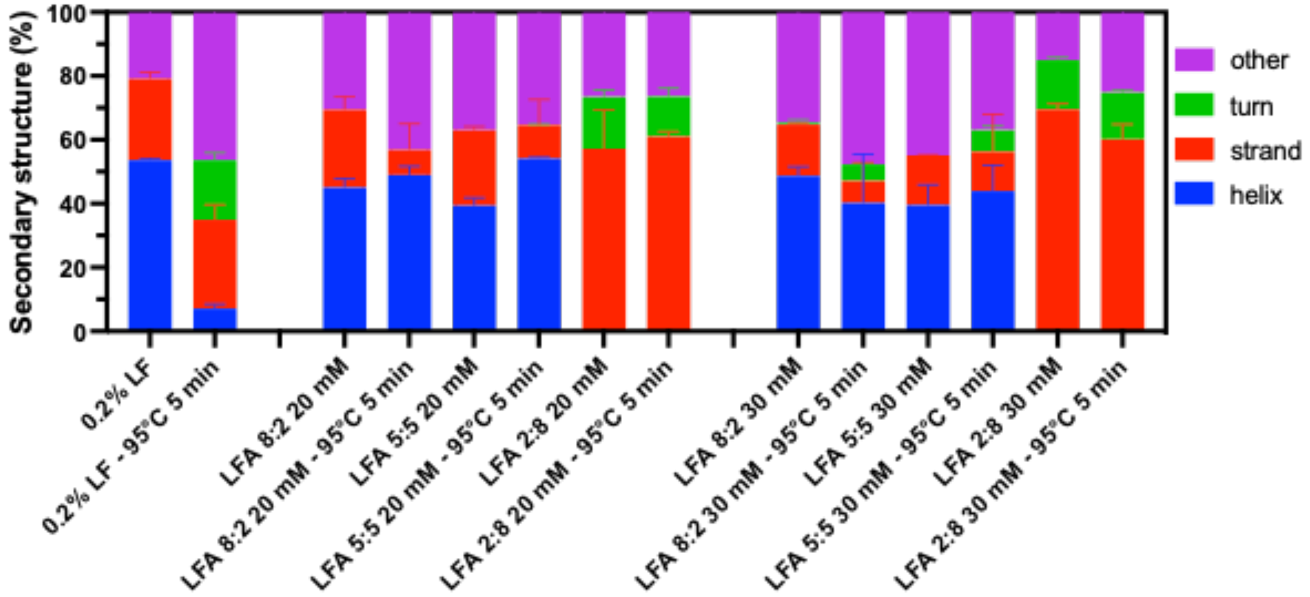


Fig. S12. The secondary structure analysis of protein and LAF hydrogels before and after heating at different LF-Alg ratios and Fe concentrations (B: LF and LF+Fe control: C: Fe 20 mM; C: Fe 30 mM).
